# Detection of astrocyte epigenetic memory in *in vitro* systems, experimental autoimmune encephalomyelitis and multiple sclerosis samples

**DOI:** 10.1101/2025.05.30.657119

**Authors:** Hong-Gyun Lee, Zhaorong Li, Camilo Faust Akl, Joon-Hyuk Lee, Gavin Piester, Jack P. Antel, Veit Rothhammer, Michael A. Wheeler, Alexandre Prat, Iain C. Clark, Francisco J. Quintana

## Abstract

We recently described astrocyte pro-inflammatory epigenetic memory based on multiple complementary *in vivo* and *in vitro* studies, and the analysis of multiple sclerosis samples. Based on bioinformatic analyses, O’Dea and Liddelow argued that the astrocyte epigenetic memory we described is the result of contamination with immune cells, particularly myeloid cells. We rebut O’Dea and Liddelow arguments as follows: **(1)** We show substantial purity of astrocytes analyzed in *in vivo* and *in vitro* systems; **(2)** We recapitulate astrocyte memory responses using five independent pure astrocyte *in vitro* systems, and show its dependency on the histone acetyl transferase p300; and **(3)** Using the Liddelow lab bioinformatic pipeline to implement purity and cell-quality criteria, we detect astrocyte epigenetic memory in five independent scRNA-seq experimental autoimmune encephalomyelitis (EAE) and multiple sclerosis (MS) astrocyte datasets. These additional analyses and studies provide further support for the existence of astrocyte pro-inflammatory epigenetic memory.

## Introduction

Astrocytes are abundant non-hematopoietic glial cells of the CNS with important functions in health and disease^1-4^. Indeed, astrocyte subsets (comprising astrocyte activation states and/or subpopulations) induced by pro-inflammatory stimuli contribute to the pathology of neurologic disorders such as MS and its model experimental autoimmune encephalomyelitis (EAE)^2,5-13^. However, the stability of astrocyte subsets, and whether they display subsequent altered responses following an initial pro-inflammatory stimulation remains unclear.

Immunological memory, the generation of faster and stronger responses upon re-stimulation, is a hallmark of T and B cell responses, driven by long-lived antigen-reactive clones^14^. Innate immune cells also display a type of immunological memory upon re-stimulation, which has been termed “trained immunity” or “inflammatory memory”^15-22^. Inflammatory memory is also reported in epithelial cells and other non-immune cell types^19,23-25^. Inflammatory memory is controlled by epigenetic writers, readers and erasers which add, recognize or remove, respectively, histone post-translational modifications (PTMs) such as acetylation or methylation. Inflammatory memory is initiated by transcription factors that, upon activation, bind specific genes, recruiting epigenetic writers that introduce histone PTMs. These PTMs alter histone interactions with DNA, increasing chromatin accessibility and consequently access to regulatory DNA sequences upon re-stimulation^26^. In addition, epigenetic readers recognize specific histone PTMs and recruit additional proteins to further modify chromatin structure and gene transcription, enabling some transcription factors to remain bound to DNA^27^. Finally, epigenetic erasers terminate inflammatory memory by removing histone PTMs^26^. However, very little is known about inflammatory memory in non-immune cells such as astrocytes.

We recently described astrocyte pro-inflammatory epigenetic memory based on multiple complementary *in vivo* and *in vitro* studies, and the analysis of MS samples^28^. Based on bioinformatic analyses, O’Dea and Liddelow argue that the astrocyte epigenetic memory that we described in animal models, *in vitro* systems, and MS samples is the result of contamination with immune cells^29^. We rebut O’Dea and Liddelow arguments as follows: **(1)** We show substantial purity of astrocytes analyzed in *in vivo* and *in vitro* systems; **(2)** We recapitulate astrocyte memory responses using five independent pure astrocyte *in vitro* systems, and show its dependency on the histone acetyl transferase p300; and **(3)** Using the Liddelow lab bioinformatic pipeline to implement purity and cell-quality criteria, we detect astrocyte epigenetic memory in five independent scRNA-seq EAE and MS astrocyte datasets. These analyses and studies confirm and further support the existence of astrocyte pro-inflammatory epigenetic memory.

## Results

### No immune cell contamination is detected in post-purification analyses of astrocyte fractions

O’Dea and Liddelow question the flow cytometry negative selection approach that we used to generate the transcriptional signature of astrocyte epigenetic memory (**Figs. 1a-d** in ^29^), suggesting that this approach results in significant immune cell contamination. However, the flow cytometry negative selection approach we used selects against multiple immune cell markers including CD11b, CD11c, CD45 and Ly6C^7,30^. Indeed, in post-purification analyses of the astrocyte fraction from our *in vivo* studies in Lee *et al.*^28^ we detected no immune cell contamination (0% CD45+, CD11b+ cells) and 93.2% GFAP+ cells (**Figs. 1a-c**). Hence, we used substantially pure astrocyte populations to generate the data in **Figs. 1a-g** in Lee *et al.*^28^. Negative selection is commonly used in these instances because it enables the active removal of contaminating immune cells, while preventing unwanted activation of cells of interest by the engagement of surface markers by soluble or plate bound antibodies. In addition, this negative selection approach diminishes the risk of selecting for specific cell subsets expressing higher levels of surface markers targeted in positive selection strategies.

**Figure 1.**
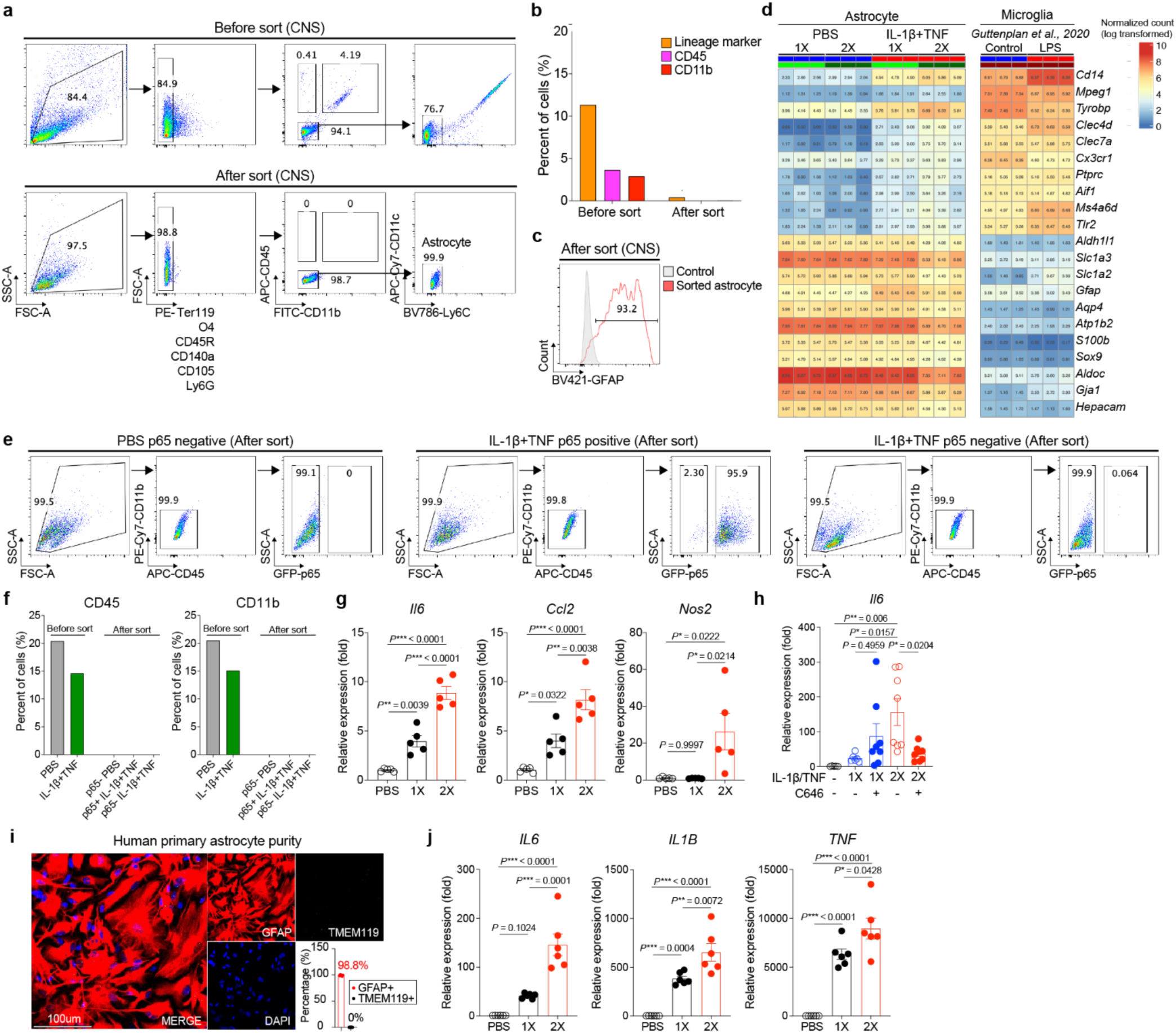
Astrocyte memory phenotype in *in vivo* and *in vitro* pure systems. **(a-c)** Fluorescence-activated cell sorting (FACS) purity analysis of mouse astrocytes before and after purification by excluding lineage-positive (Ter119, O4, CD45R, CD140a, CD105, Ly6G) and myeloid/lymphocyte-positive (CD45, CD11b, CD11c, Ly6C) cells. **(d)** Differential gene expression determined by RNA-seq in FACS-sorted astrocytes from C57BL/6 mice following intracerebroventricular (ICV) administration of IL-1β and TNF or PBS once (1X) or twice (2X) (left panel)^28^. Differential gene expression determined by RNA-seq in MACS purified microglia stimulated with/without LPS (right panel)^38^. **(e-f)** Purity analysis of primary astrocytes used in Figure 1i in Lee et al^28^. In vitro GFP+ and GFP− astrocyte fractions were isolated by FACS excluding myeloid-positive (CD45, CD11b) cells from mouse p65-eGFP reporter astrocytes following 18-hour stimulation with PBS or IL-1β + TNF. **(g)** Quantitative PCR (qPCR) of CD11b-CD45-O4-negative FACS-sorted astrocytes after 30min stimulation with IL-1β/TNF stimulation once (1X) or twice (2X) (n=5 per group). One-way ANOVA followed by Tukey’s multiple comparison test. **(h)** Immunopanned primary astrocytes received IL-1β/TNF or control stimulation in the presence or absence of C646 (p300/CBP inhibitor). qPCR performed after 1h stimulation with IL-1β/TNF on day 7 (n=5-8 per group). One-way ANOVA followed by Tukey’s multiple comparison test. **(i)** Purity analysis of human primary astrocytes used in Extended Data Figure 9a in Lee et al^28^. **(j)** qPCR of human iPSC-derived astrocytes that received IL-1β/TNF stimulation once (1X) or twice (2X) (n=6 per group). One-way ANOVA followed by Tukey’s multiple comparison test.

### Purified astrocytes express canonical markers

O’Dea and Liddelow raise concerns about the transcriptional signature of astrocyte memory, arguing that astrocytes purified by negative selection against immune cell markers and stimulated twice with IL-1β+TNF show a relative decrease in astrocyte marker expression relative to astrocytes stimulated only once (**Fig. 1a** in ^29^). However, although the expression of astrocyte markers is slightly reduced by 2x IL-1β+TNF stimulation (not by 1x IL-1β+TNF condition), their absolute expression levels remain high, identifying these cells as astrocytes **(Fig. 1d)**. These findings agree with reports of the decreased expression of some astrocyte canonical markers such as *Aqp4, Slc1a2, Slc1a3* and *Hepacam* in response to inflammatory stimuli^31^. Indeed, the Liddelow lab reported the downregulation of GJA1 in response to inflammatory challenge^32^.

### Astrocytes express non-canonical genes in the context of inflammation

O’Dea and Liddelow raise concern by the detection in astrocytes stimulated twice with IL-1β+TNF of some genes also expressed by immune cells (**Fig. 1a** in ^29^). However, the expression of these genes is low compared to the expression of astrocyte markers, or to their own expression in microglia (**Fig. 1d**). O’Dea and Liddelow frequently represent our data using normalized z-scoring, which distorts the display of canonical and non-canonical genes expressed by astrocytes^29^. As an example, we show raw counts versus z-score normalized marker expression in astrocytes from the Allen Atlas dataset; the raw counts demonstrate high astrocyte marker expression while z-scoring makes many cell clusters appear as if they do not include astrocytes **(Extended Data Figs. 1a-b)**.

In addition, the expression in astrocytes of genes expressed by hematopoietic cells has been reported by many other groups^10,13,31-36^. For instance, the canonical monocyte marker *Cd14* was identified by the Barres lab as a marker of A2 reactive astrocytes^10,34^. Additional markers highlighted by O’Dea and Liddelow have been detected in astrocytes including *S100a8* in sepsis^35^ and *Tyrobp* in multiple disease models^13,31,36^. *Cd74* and *H2ab1* were detected in astrocytes^31^ and recently highlighted in a review by the Liddelow group^37^. Indeed, datasets generated by the Liddelow group^33,38^ (**Extended Data Figs. 1c-d**) and others including the Allen Brain Cell dataset^31,36,39-42^ (**Extended Data Figs. 1e-j**) show expression in astrocytes of myeloid/lymphoid cell markers (highlighted in red), and the downregulation of astrocyte markers (highlighted in blue) upon inflammatory stimulation (**Extended Data Figs. 1c-j**). Of particular note is the detection of genes associated with other cell types in astrocytes purified by the Liddelow lab by immunopanning (**Extended Data Fig. 1c**)^38^, an approach suggested by Liddelow and O’Dea to study astrocyte epigenetic memory without concerns of immune cell contamination^29^.

In summary, the transcriptional downregulation of homeostatic astrocyte markers and the upregulation of some genes linked to other cell types is a feature of reactive astrocytes widely reported by other groups including Liddelow’s, recapitulating similar observations made in other cell types such as neurons^43^, oligodendrocytes^44^ and even fibroblasts^24,45^ in the context of inflammation and pathology.

### The use of the Allen Brain Cell dataset to study astrocytes in inflammation is misleading

O’Dea and Liddelow examined the transcriptional signature of astrocyte memory using transcriptional modules linked to cell types in the Allen Brain Cell atlas^42^; they state that this analysis detects immune cell contamination (**Figs. 1b-d** in ^29^). However, analyzing astrocytes under pro-inflammatory conditions using the Allen dataset built under homeostatic conditions is a biased approach as acknowledged by O’Dea and Liddelow themselves^29^: “*The Allen dataset is from wild type mouse brain without an inflammatory insult, and it is possible that some of the up-signature genes are restricted to immune cells in the healthy brain but induced in astrocytes during inflammation*”.

To exemplify how misleading this approach as applied by O’Dea and Liddelow^29^ is, we used the Allen dataset to analyze the response to activation detected by single cell or bulk RNA-seq of primary astrocytes generated by the Liddelow lab according their criteria of purity and quality^33,38^. Using the Allen dataset to analyze the transcriptional response of astrocytes purified by immunopanning and stimulated *in vitro* with IL-1α+TNF+C1q- or *in vivo* with LPS reported by Liddelow and coworkers^33,38^, we detect an upregulation of transcriptional modules linked to “contaminating immune cells” (e.g., monocytes) as referred to by O’Dea and Liddelow **(Extended Data Figs. 2a-b**). In addition, we identify the transcriptional module linked to astrocytes amongst the most downregulated ones **(Extended Data Figs. 2a-b**). We obtained similar results when we used the Allen dataset to analyze additional datasets of the transcriptional response of astrocytes generated by multiple independent groups^31,36,39,41,46-48^; these analyses revealed an upregulation of transcriptional modules linked to other cell types and a downregulation of those linked to astrocytes **(Extended Data Figs. 2c-i)**. Collectively, these findings demonstrate that the use of the Allen dataset built on naïve conditions to analyze astrocytes in the context of pathology as done by O’Dea and Liddelow is misleading.

### Astrocyte epigenetic memory is detected in five independent pure in vitro astrocyte systems

O’Dea and Liddelow suggest the use *in vitro* pure astrocyte systems to demonstrate that astrocyte epigenetic memory is not driven by immune cell contamination^29^. Indeed, to focus on the contribution of astrocyte intrinsic mechanisms of astrocyte pro-inflammatory memory, we used multiple independent *in vitro* systems to study astrocyte-intrinsic epigenetic memory independently of the trained responses of other cells in the CNS^28^.

In one of these systems, we prepared cultures of NF-κB reporter (p65-GFP) astrocytes as done for decades in the astrocyte field^49-90^, and activated them with the known NF-κB activators IL-1β and TNF (IL-1β+TNF). Following this first *in vitro* stimulation, we sorted by flow cytometry p65-GFP+ and p65-GFP-astrocytes selectively excluding CD45-positive and CD11b-positive cells; no immune cell contamination (0% CD45+ or CD11b+ cells) is detected in post-sorting analyses of the p65-GFP+ astrocyte fraction **(Figs. 1e-f)**. p65-GFP+ and p65-GFP-astrocytes were then rested for a week and re-stimulated with IL-1β+TNF, detecting stronger responses in the previously stimulated p65-GFP+ fraction **(Fig. 1i and Extended Data Fig. 1m** in Lee *et al.*^28^), supporting the existence of astrocyte inflammatory memory. Since the p65-GFP+ astrocyte fraction shows no immune cell contamination (0% CD45+ or CD11b+ cells) after sorting, before resting and restimulation **(Figs. 1e-f)**, this demonstrates that the memory phenotype detected upon re-stimulation is driven by astrocytes and not contaminating immune cells. O’Dea and Liddelow claim 15% contamination by other glial cells after sorting p65-GFP+ astrocytes^29^, although this point was clarified both in our manuscript **(Fig. 1i and Extended Data Fig. 1m** in Lee *et al.*^28^).

To further study memory in pure astrocytes, we eliminated non-astrocyte populations by flow cytometry sorting prior to the initiation of mouse primary astrocyte cultures, before the first *in vitro* IL-1β+TNF stimulation. Specifically, we negatively selected against CD11b⁺, CD45⁺ and Olig4⁺ cells to remove microglia, immune cells, and oligodendrocytes, respectively (**Extended Data Figs. 3a-b**). We then stimulated astrocytes twice with IL-1β+TNF, one week apart. Even under this stringent purification, we detected a robust memory response upon re-stimulation (**Fig. 1g**), further confirming the existence of astrocyte memory in the absence of contaminating immune cells.

In addition, we also detected astrocyte memory in pure *in vitro* astrocyte cultures established by immunopanning **(Fig. 1h, Extended Data Fig. 3c)**, a system suggested by O’Dea and Liddelow to study astrocyte-intrinsic epigenetic memory in the absence of immune cell contamination^29^.

Finally, we also demonstrated the induction of pro-inflammatory memory *in vitro* in human primary astrocytes (**Extended Data Fig. 9a** in Lee *et al.*^28^) devoid of contaminating microglia as confirmed by immunofluorescence (**Fig. 1i**). We further validated these findings using microglia-free human induced pluripotent stem cell (iPSC)-derived astrocytes (**Fig. 1j, Extended Data Figs. 3d-e**) as suggested by O’Dea and Liddelow^29^. Indeed, it is surprising that O’Dea and Liddelow suggest the use of this system to validate the existence of astrocyte memory^29^, when we shared by email these iPSC-derived astrocyte data with them months before the upload of their piece to BioRxiv.

In summary, we detect astrocyte inflammatory memory in five independent mouse and human pure astrocyte systems. However, please note that in the **Discussion** of Lee *et al.*^28^ we postulate that other cells such as microglia, contribute to astrocyte inflammatory memory *in vivo*: “*Indeed, although we focused on astrocyte-intrinsic mechanisms of epigenetic memory, interactions with other cells in the CNS such as microglia are also likely to modulate astrocyte epigenetic memory*”.

### Astrocyte epigenetic memory is dependent on p300

O’Dea and Liddelow argue that we provided insufficient evidence that p300 participates in astrocyte epigenetic memory^29^. However, our conclusions on the role of p300 in astrocyte epigenetic memory were based on complementary *in vivo* and *in vitro* studies as follows:

**First**, we inactivated *Ep300* in astrocytes *in vivo* by inducing frameshift mutations using a lentivirus that expresses sgRNA and CRISPR/Cas9 under the control of the *GfaABC_1_D* promoter. O’Dea and Liddelow point out that *Ep300* mRNA levels appeared unchanged in our RNA-seq analysis (**Figs. 1e-f** in ^29^). However, this observation is compatible with the introduction of frameshift mutations which alter protein translation but not mRNA transcription^91^. This is also a common observation particularly when using relatively low-depth RNA-seq, which has limited sensitivity to detect changes in moderately or lowly expressed genes^92-94^. In fact, CRISPR-based perturbation studies -including those targeting *Ep300*-have reported no reduction in mRNA levels despite protein depletion^91,95-99^. Most importantly, we validated the loss of p300 protein expression (and function, see below) induced by CRISPR/Cas9-induced frameshift mutations (**Extended Data Figs. 3f, 7a** in Lee *et al.*^28^).

**Second,** we showed that *Ep300* inactivation in astrocytes reduces histone acetylation (H3K27ac), a product of p300 histone acetyltransferase activity, in genes related to inflammation as determined by chromatin immunoprecipitation followed by sequencing (ChIP-seq) (**Figs. 2e-g**, in Lee *et al.*^28^).

**Figure 2.**
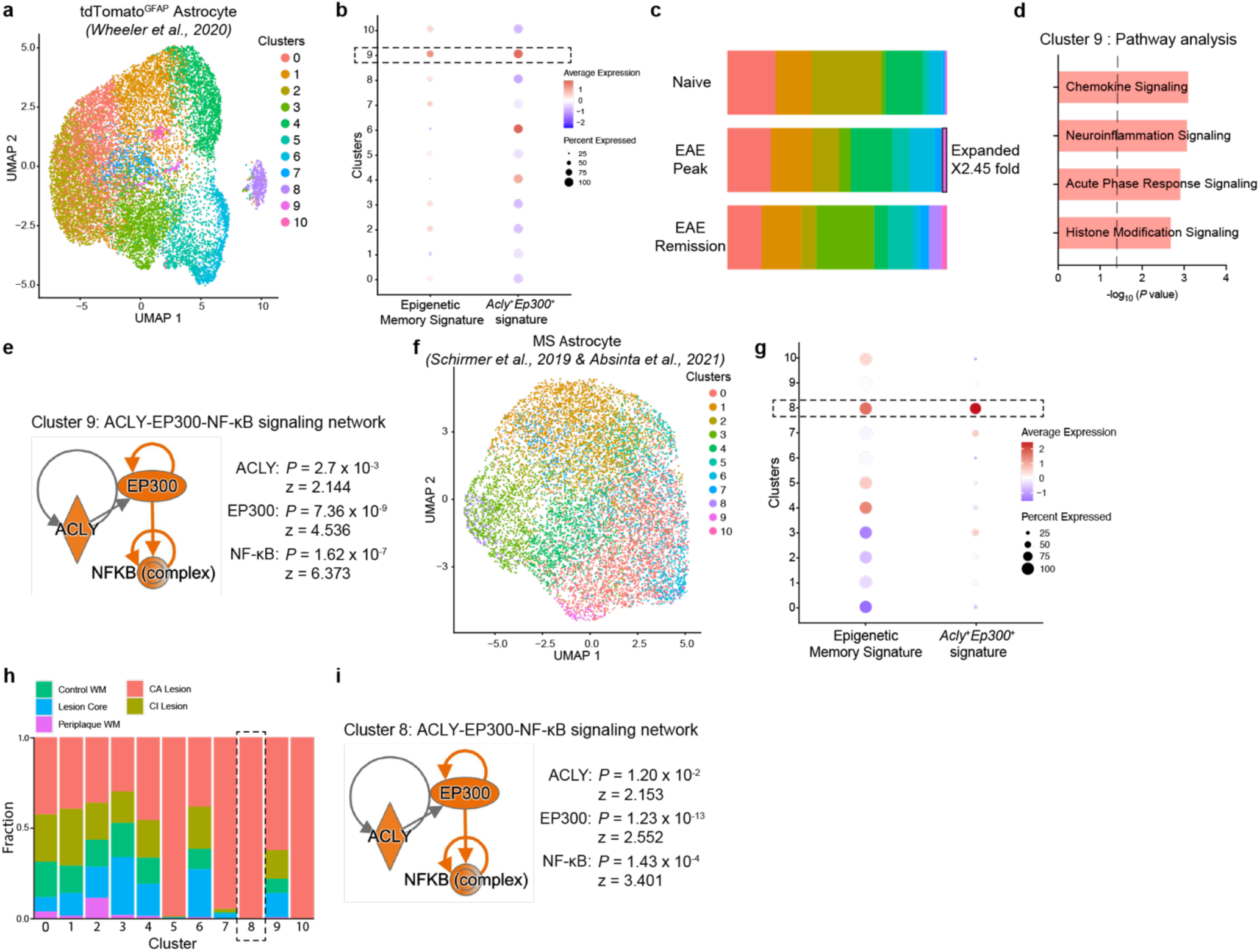
Detecton of memory astrocyte subsets in EAE and MS using Liddelow lab pipeline. **(a)** UMAP plot of TdTomato^Gfap^ astrocytes applying Hasel *et al.^33^* filtering criteria. **(b)** *Acly*^+^*Ep300*^+^ EAE astrocyte signature and astrocyte epigenetic memory signature expression in EAE astrocyte clusters. **(c)** Fraction of cells per cluster overall. **(d)** IPA up-regulated pathways (red) in cluster 9 astrocytes. **(e)** ACLY–EP300–NF-κB signaling network in cluster 9 astrocytes. **(f)** UMAP plot of astrocytes from MS and control samples from Schirmer *et al.*^109^ and Absinta *et al.*^110^ applying Hasel *et al.^33^* filtering criteria. **(g)** *Acly*^+^*Ep300*^+^ EAE astrocyte signature and astrocyte epigenetic memory signature expression in MS astrocyte clusters. **(h)** Astrocyte cluster composition analysis in MS. WM, white matter; CI, chronic inactive; CA, chronic active. **(i)** ACLY–EP300–NF-κB signaling network of cluster 8 astrocytes.

**Third,** we used gene set enrichment analysis (GSEA) to establish that *Ep300* inactivation in astrocytes significantly decreases the astrocyte epigenetic memory up-signature in EAE (**Fig. 3c, Extended Data Fig. 6h** in Lee *et al.*^28^). In addition, our GSEA core enrichment analysis revealed that p300 knockdown in astrocytes significantly reduced the expression of 12% (11 out of 91) and 19.7% (18 out of 91) of genes within the epigenetic memory astrocyte up-signature in B6-EAE and NOD-EAE, respectively; additional epigenetic modifiers likely participate in the control of astrocyte epigenetic memory as we suggested (see below). GSEA is widely used in transcriptomic studies^100^ because it evaluates changes at the pathway level rather than relying solely on large changes on the expression of individual genes, an important point because the effects of epigenetic regulators on gene expression are frequently studied at the pathway level while considering gene-gene correlations^101-104^. Indeed, GSEA was developed to detect alterations in pathway activity based on the analysis of the relative rank of multiple co-regulated genes in a dataset, meaning that also non-statistically significant changes in the expression of all individual genes in a pathway are analyzed in parallel to detect significant effects on pathway activity. Thus, since GSEA is designed to analyze unfiltered ranked lists of all genes in a pathway, we analyzed the effect of *Ep300* inactivation in astrocytes on all genes in the memory astrocyte up-signature preranked based on log2-fold-change, in line with the <u>GSEA User Guide</u>: *“**Filtering based on expression values.** …The GSEA algorithm does not filter the expression dataset and generally does not benefit from your filtering of the expression dataset. During the analysis, genes that are poorly expressed or that have low variance across the dataset populate the middle of the ranked gene list and the use of a weighted statistic ensures that they do not contribute to a positive enrichment score. By removing such genes from your dataset, you may actually reduce the power of the statistic…”*.

**Fourth,** in complementary studies using our *in vitro* memory model in mouse primary astrocyte cultures to address limitations of *in vivo* studies (*e.g.*, concomitant training of immune cells) we detected a significant increase in p300 recruitment to the promoters of pro-inflammatory genes in astrocytes, including *Il6*, *Ccl2*, and *Nos2* (**Extended Data Fig. 3m** in Lee *et al.*^28^).

**Fifth,** pharmacological inhibition of p300 using C646 suppressed the induction of astrocyte inflammatory memory *in vitro* in both human and mouse microglia-free astrocyte cultures **(Figs. 1e-f, h,i Extended Data Fig. 3c** in this article; **Fig. 1j and Extended Data Fig., 9a** in Lee *et al.*^28^), further demonstrating the need of p300 activity for the induction of astrocyte memory. In one of these studies we treated p65-GFP reporter astrocytes with C646 during the first IL-1β+TNF stimulation, sorted GFP+ astrocytes, rested them and evaluated their response to IL-1β+TNF re-stimulation in the absence of the “immune cell contamination” that O’Dea and Liddelow claim is the source of astrocyte memory **(Fig. 1j and Extended Data Fig. 1m** in Lee *et al.*^28^). Similarly, inhibition of p300 also prevented the memory response to IL-1β+TNF restimulation (but not the response to first IL-1β+TNF stimulation) in mouse astrocytes purified by immunopanning, a system suggested by O’Dea and Liddelow to demonstrate the existence of astrocyte-intrinsic memory in the absence of immune cell contamination^29^ **(Fig. 1h, Extended Data Fig. 3c)**.

In summary, these findings provide robust, complementary *in vivo* and *in vitro* evidence that p300 participates in the control of astrocyte epigenetic memory. Additional epigenetic modifications and modifiers likely contribute to astrocyte epigenetic memory, as we mentioned in our manuscript^28^.

Finally, O’Dea and Liddelow argue that because p300 and ACLY expression is not restricted to astrocytes (**Fig. 1g** in ^29^), they do not define an epigenetic memory state specific to astrocytes. In our manuscript we monitor p300 and ACLY expression in the context of their co-expression with astrocyte markers or in purified astrocytes in EAE, MS or *in vitro* systems (**Figs. 3g, 5i and Extended Data Figs. 2b, 4a, 5f, 5i, 6e**, **10a** in Lee *et al.*^28^). In addition, in the **Discussion** section we highlight that shared mechanisms regulate the epigenetic state of multiple cell types; referencing other cell types where p300 also regulates similar epigenetic memory responses: *“The brain displays the highest p300/CBP HAT activity in the body. Indeed, histone acetylation and other post-translational modifications, have been linked to long-term memory including recognition and contextual fear. p300 also regulates cellular senescence and stress response, which promote systemic chronic inflammation.”* However, our ATAC-seq, ChIP-seq, and ChIP-qPCR data demonstrate that p300 and ACLY participate in the regulation of astrocyte epigenetic memory in inflammatory conditions **(Figs. 1f-g, 2e-g, Extended Data Figs. 1d-f, 3l-m** in Lee *et al.*^28^). Moreover, our *in vitro* astrocyte culture and *in vivo* immunostaining data using astrocyte markers show p300 and ACLY upregulation in astrocytes in B6-EAE (**Fig. 3g, Extended Data Fig. 5f in** Lee *et al.*^28^), NOD-EAE (**Extended Data Fig. 6e** in Lee *et al.*^28^), MS lesions (**Fig. 5i, Extended Data Fig. 10a** in Lee *et al.*^28^), and inflammatory conditions **(Extended Data Figs. 2b, 4a** in Lee *et al.*^28^). Similarly, Sofroniew and colleagues reported p300 upregulation in astrocytes following LPS treatment or spinal cord injury^36^. Finally, we showed that *Ep300* and *Acly* inactivation in astrocytes reduces histone acetylation H3K27ac and impairs the astrocyte epigenetic memory signature in B6-EAE **(Figs. 2d, 3c, Extended Data Figs. 3f, 5a** in Lee *et al.*^28^) and NOD-EAE **(Extended Data Figs. 6h, 7a-b** in Lee *et al.*^28^). Thus, based on multiple complementary approaches we conclude that p300 and ACLY participate in the control of astrocyte epigenetic memory, while stating in the **Discussion** that additional epigenetic mechanisms (e.g. histone serotonylation) likely participate in this process.

### Memory astrocyte detection in 3 independent EAE studies using the Liddelow lab filtering criteria

O’Dea and Liddelow raise two main criticisms regarding the identification of memory astrocytes in EAE by scRNA-seq. First, they argue that astrocytes should not express genes linked to other cell types. Second, they claim that the scRNA-seq dataset of TdTomato+ astrocytes in EAE (*Gfap^TdTomato^* mice)^105,106^ is contaminated and memory astrocytes are not detected in a dataset of “pure” astrocytes, as per definition.

As addressed above, the first point is not supported by multiple reports^31,36,39-41^—including from the Liddelow lab*^33,38^*—which show the expression in astrocytes of most, if not all, of the genes highlighted by O’Dea and Liddelow in their critique **(Extended Data Figs. 1c-j**). Since these genes are detected by bulk RNA-seq and spatial transcriptomics, indeed they will be likely detected by scRNA-seq in individual cells. However, the expression of these genes in TdTomato+ cells is much lower than the expression of astrocyte markers when samples are analyzed in pseudobulk **(Extended Data Fig. 4a)**.

To address the second point, we reanalyzed our *Gfap^TdTomato^*astrocyte dataset according to the same criteria used by the Liddelow lab to reanalyze our *Gfap^TdTomato^* dataset for their Hasel *et al.* publications^33,107^. When the Liddelow lab analyzed our *Gfap^TdTomato^* dataset, it integrated seamlessly with their own dataset of *Aldh1l1::EGFP* astrocytes (see **Figs. 7e-f** in ^33^). Indeed, in their previous studies over half of our tdTomato dataset was seamlessly incorporated into their own astrocyte scRNA-seq analyses^33,107^. However, the CellTypist analysis they now provide suggests that only cluster 0 from our *Gfap^TdTomato^* dataset contains astrocytes (**Fig. 2e** in^29^), contradicting theor previous integration and use of this dataset^33,107^.

Using the Liddelow lab’s code on GitHub, we reanalyzed our *Gfap^TdTomato^*dataset by selecting clusters enriched for astrocyte marker genes (e.g., *Aldoc*, *Aqp4*, *Slc1a3*, *Slc1a2*) and de-enriched for genes associated with myeloid, endothelial, neuron, and oligodendrocyte lineages. We retained 4 clusters from this analysis, which is the same number of clusters retained when this dataset was reanalyzed by the Liddelow lab^33^, and performed astrocyte re-clustering. Please note that the cells selected in this analysis robustly express astrocyte marker genes and virtually none of the genes O’Dea and Liddelow note as concerning **(Extended Data Fig. 4b)**. We then examined the transcriptional memory signature in this dataset. In cluster 9, which is expanded in peak EAE, we detected significant induction of both the *Acly*+*Ep300*+ FIND-seq signature and the epigenetic memory signature (**Figs. 2a-e**). Thus, we still detect the memory phenotype when we apply the bioinformatic pipeline from the Liddelow lab.

We shared the above analysis with O’Dea and Liddelow before the upload of their BioRxiv article, but they do not acknowledge the existence of this re-analysis in their critique. However, they do acknowledge it in their Jupyter notebooks (https://github.com/michael-r-odea/EpiMemAstros), where they do not disagree about the purity of the re-analyzed samples. They now argue that this re-analysis is insufficient because we removed most “memory cells” when filtering to address their original criticisms. To address this concern, we used the same Liddelow lab^33^ criteria to re-analyze two independent^47,48^ EAE scRNA-seq datasets, and we still detect a cluster of astrocytes displaying the transcriptional signature linked to epigenetic memory **(Extended Data Figs. 5a-j)**; this cluster is expanded in EAE in both datasets. In addition, O’Dea and Liddelow now claim that because the astrocyte memory cluster is less numerous, it has no physiologic relevance. This argument is inconsistent with publications from the Liddelow lab describing IFN-responsive and *Myoc*+ astrocyte subsets encompassing as few as 2.7%^33^ and 2.04%^107^ astrocytes, which they claim are physiologically relevant without any functional validation. Most importantly, a rare glutamatergic astrocyte subset was recently shown to play a critical role in learning and memory^108^, highlighting the physiological relevance of small astrocyte subsets.

In summary, using the Liddelow’s lab bioinformatic pipeline, we detect astrocytes displaying a transcriptional signature associated with epigenetic memory in our EAE TdTomato+ astrocyte scRNA-seq dataset and in two independent EAE scRNA-seq datasets generated by other investigators^47,48^.

### Memory astrocyte detection in 2 MS snRNA-seq datasets using the Liddelow lab filtering criteria

O’Dea and Liddelow criticized our re-analysis of human MS astrocyte snRNA-seq datasets generated by Arnold Kriegstein and David Rowitch^109^, and Peter Calabresi and Daniel Reich^110^. We reanalyzed these datasets because they enabled the study of similar lesion types and brain areas after undergoing substantial quality control filtering and cell type annotation at the time of their publication^109,110^. Specifically, we extracted astrocytes as identified on the metadata from processed snRNA-seq objects, meaning that the astrocytes analyzed in Lee *et al.*^28^ were those already identified and validated as astrocytes by these independent groups^109,110^. Of note, extracting cell types based on pre-existing metadata was sufficient for O’Dea and Liddelow when they performed their Allen Brain Cell Atlas analyses, using astrocytes where some myeloid gene expression is detected **(Extended Data Fig. 1j)**. However, they criticize this strategy when we use it to analyze other datasets. Nevertheless, we validated astrocyte marker expression by the Schirmer *et al.* dataset in **Extended Data Figs. 9a, 10a** in Wheeler *et al.*^105^, and successfully clustered the Schirmer *et al.* data with 3 other astrocyte scRNA-seq datasets^105^, indicating they are consistent with astrocyte data from other groups. The Absinta *et al.* dataset^110^ was released in 2021, after Wheeler *et al.* was published^105^.

O’Dea and Liddelow claim the Absinta *et al.*^110^ and Schirmer *et al.*^109^ datasets are contaminated with non-astrocyte populations and low quality cells, invalidating our analysis. However, astrocytes in these datasets have been already filtered for purity and quality control^109,110^. Indeed, the use of z-scoring to display marker expression in the absence of other cell types is misleading (Fig. 2d, 3i in ^29^), as it suggests lower expression of canonical markers than when displaying raw counts **(Extended Data Figs. 6a-b)**.

In addition, O’Dea and Liddelow argue that because the memory signatures we defined are contaminated by non-astrocytes, they should be undetectable in populations of pure astrocytes. To address these concerns, we re-analyzed these snRNA-seq datasets using parameters specified by the Liddelow lab^33^, eliminating clusters expressing the markers they criticize, as well as any cluster expressing marker genes associated with other cell populations in homeostasis. We then removed one cluster displaying overall lower UMIs and genes relative to other clusters. The cells we bioinformatically selected robustly express astrocyte marker genes **(Extended Data Fig. 4c)** and feature high-quality nuclei **(Extended Data Fig. 4d)**. In this dataset, we detected co-expression of the epigenetic memory signature and *Acly*+*Ep300*+ FIND-seq signature in cluster 8, which was expanded in MS chronic active lesions **(Figs. 2f-i)**. Thus, we detected an astrocyte subset displaying a transcriptional signature linked to epigenetic memory and associated with chronic active lesions in an astrocyte dataset devoid of the cell-type markers and low-quality cells that O’Dea and Liddelow state as being concerning.

In their Jupyter notebooks (https://github.com/michael-r-odea/EpiMemAstros) O’Dea and Liddelow critique the number of cells exhibiting the memory signature, but not the purity of these samples. Please note that we detect the transcriptional signature of astrocyte memory in active lesions, which are relatively small regions of MS pathology. Moreover, the critiques of O’Dea and Liddelow surrounding abundance are not in agreement with their own publications, given their claims of physiological roles for astrocyte populations of ∼2% abundance^33,107^. Most importantly, we validated by immunostaining in independent samples of MS active lesions the increase in ACLY+p300+ astrocytes **(Fig. 5i, Extended Data Fig. 10a** in Lee *et al.*^28^).

As with our re-analysis of the TdTomato+ cell data above, O’Dea and Liddelow claim that since we removed the original “memory astrocytes” they critiqued, our paradigm is invalid when the signature is applied to filtered astrocytes. Thus, to further strengthen our findings, we analyzed an additional MS scRNA-seq dataset^111^, detecting an astrocyte subset expanded in MS displaying a transcriptional signature linked to memory **(Extended Data Figs. 7a-e)**.

In summary, we reproduced our initial findings on EAE and MS by re-analyzing our original datasets using Liddelow’s bioinformatic pipelines, and by analyzing independent datasets generated by other laboratories. Since the Liddelow lab pipeline discards astrocytes expressing mRNAs linked to other cell types, even when other labs showed these genes are expressed in astrocytes, this re-analysis is likely to underestimate the abundance of astrocytes displaying a transcriptional signature of memory.

## Conclusion

Based on bioinformatic analyses, O’Dea and Liddelow argue that astrocyte memory reflects immune cell contamination. We refute this claim with multiple complementary experimental and bioinformatic data as follows: **1)** We show that astrocyte fractions *in vitro* and *in vivo* are free of contaminating immune cells; **2)** We show that datasets generated by multiple independent labs, including Liddelow’s, detect the downregulation of astrocyte marker expression and the upregulation of mRNAs linked to hematopoietic cells in mouse and human astrocytes in the context of inflammation; **3)** We show the inadequacy of using Allen Brain Cell Atlas transcriptional modules generated under homeostatic conditions to analyze astrocytes in the context of inflammation as done by O’Dea and Liddelow; **4)** We present new experimental data demonstrating immune memory in pure astrocyte culture systems: FACS-sorted mouse astrocytes, immunopanned mouse astrocytes, and human iPSC-derived astrocytes (**Figs. 1g-h,j)**. Importantly, we also show that astrocyte memory in immunopanned astrocytes is dependent on p300 (**Fig. 1h)**; and **5)** Using Liddelow’s bioinformatic pipelines designed to remove “contaminating and bad quality cells” we detect astrocytes displaying a transcriptional signature of memory in our EAE and MS datasets, and in independent EAE and MS datasets from other labs.

It is worth highlighting points where we agree with O’Dea and Liddelow. **First,** O’Dea and Liddelow agree that astrocyte immune memory exists and do not challenge the principle we proposed^29^. **Second,** O’Dea and Liddelow suggest the use of *in vitro* pure astrocyte systems, including immunopanning and iPSC-derived astrocytes, to study astrocyte memory in the absence of immune cell contamination^29^. All culture systems have inherent limitations, but nonetheless remain valuable tools. For example, immunopanning has demonstrated to yield lower cellular purity, but it potentially provides higher cell viability than positive selection by flow cytometry sorting^112^. Similarly, iPSC-derived astrocytes, although promising for modeling CNS diseases, still face challenges in reliably generating fully mature astrocytes representative of their counterparts in the CNS^113-118^. Nonetheless, we have now validated astrocyte memory in no less than 5 independent *in vitro* systems devoid of contaminating immune cells, including the immunopanning- and iPSC-derived astrocytes. Hence, we have proven the existence of astrocyte-intrinsic epigenetic memory by O’Dea and Liddelow’s own definitions and standards.

In summary, our multiple and complementary validation analysis and functional interrogation studies support the existence of astrocyte-intrinsic epigenetic memory independent of immune cell contamination, a phenomenon also reported by others^40,119-121^.

**Extended Data Figure 1.**
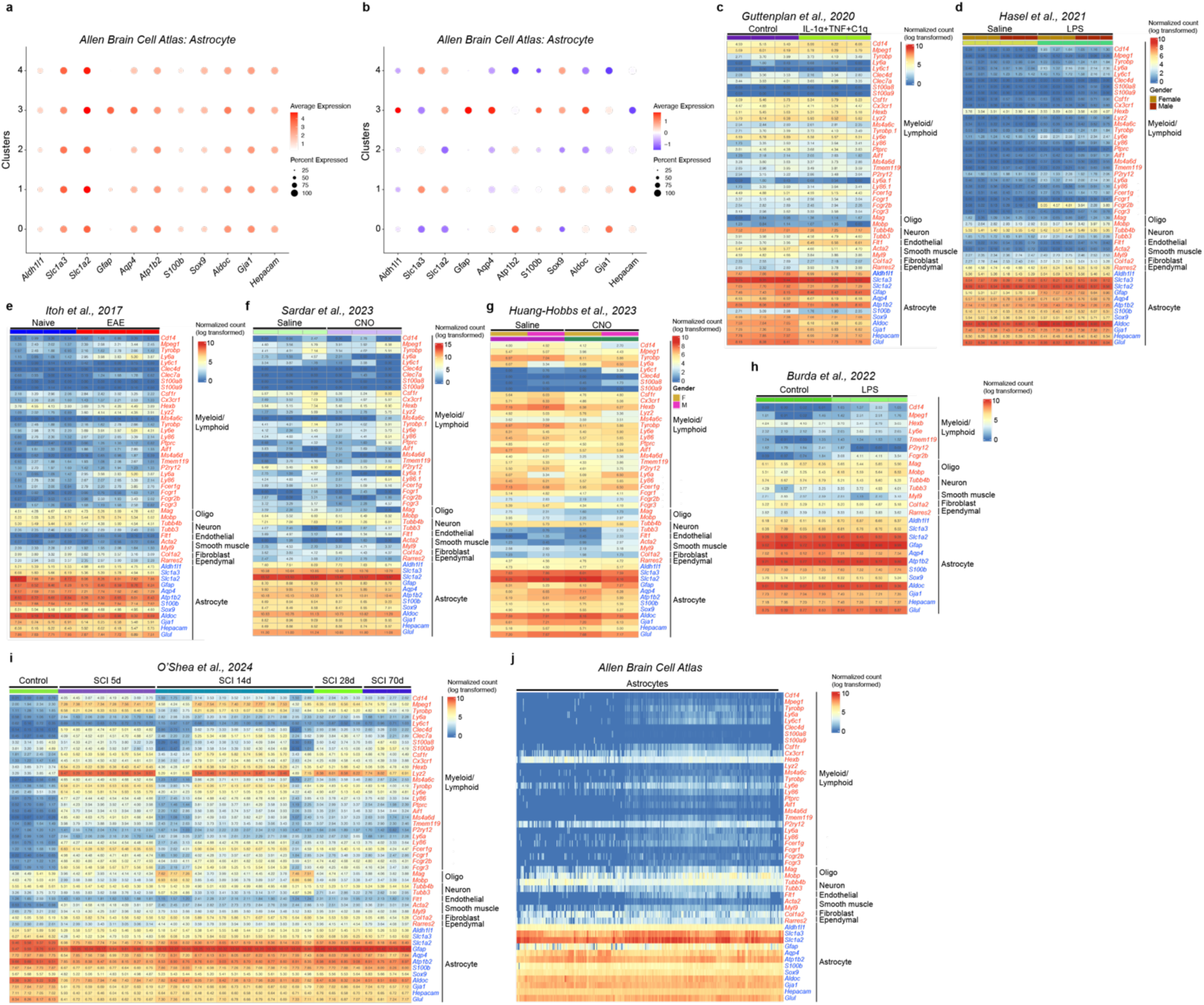
Astrocyte expression of canonical and non-canonical genes in independent datasets. **(a)** Astrocyte dot plot of canonical cell type marker genes across the clusters from Allen Brain Cell Atlas*^42^* based on normalized counts. **(b)** Astrocyte dot plot of canonical cell type marker genes across the clusters from Allen Brain Cell Atlas*^42^* based on scaled counts (subtracted by mean and divided by standard deviation). **(c)** Gene expression of immunopanned astrocytes stimulated by IL-1α, TNFα, and C1q or control determined by RNA-seq^38^. **(d)** Gene expression of FACS-sorted astrocytes from Aldh1l1^eGFP^ mice treated with saline or LPS-injected determined by single-cell RNA-seq^33^. **(e)** Gene expression in astrocytes from control or EAE astrocyte-specific ribotag mice (GFAP-Cre:RiboTag) mice determined by RNA-seq analysis^39^. **(f)** Gene expression of FACS-sorted astrocytes from Gq-CNO or Gq-Saline injected Aldh1l1^eGFP^ mice determined by RNA-seq^40^. **(g)** Gene expression of astrocytes determined by single cell RNA-seq of glioma-bearing mice treated with CNO or Saline^41^. **(h)** Gene expression determined by RNA-seq in astrocytes from astrocyte-specific Ribotag mice (GFAP-Cre:RiboTag) treated with saline or LPS^36^. **(i)** Gene expression determined by RNA-seq in astrocytes from astrocyte-specific Ribotag mice (Aldh1l1-CreERT-RiboTag)^31^ after spinal cord injury (SCI) or control treatment. **(j)** Gene expression in astrocytes from the Allen Brain Cell Atlas*^42^*.

**Extended Data Figure 2.**
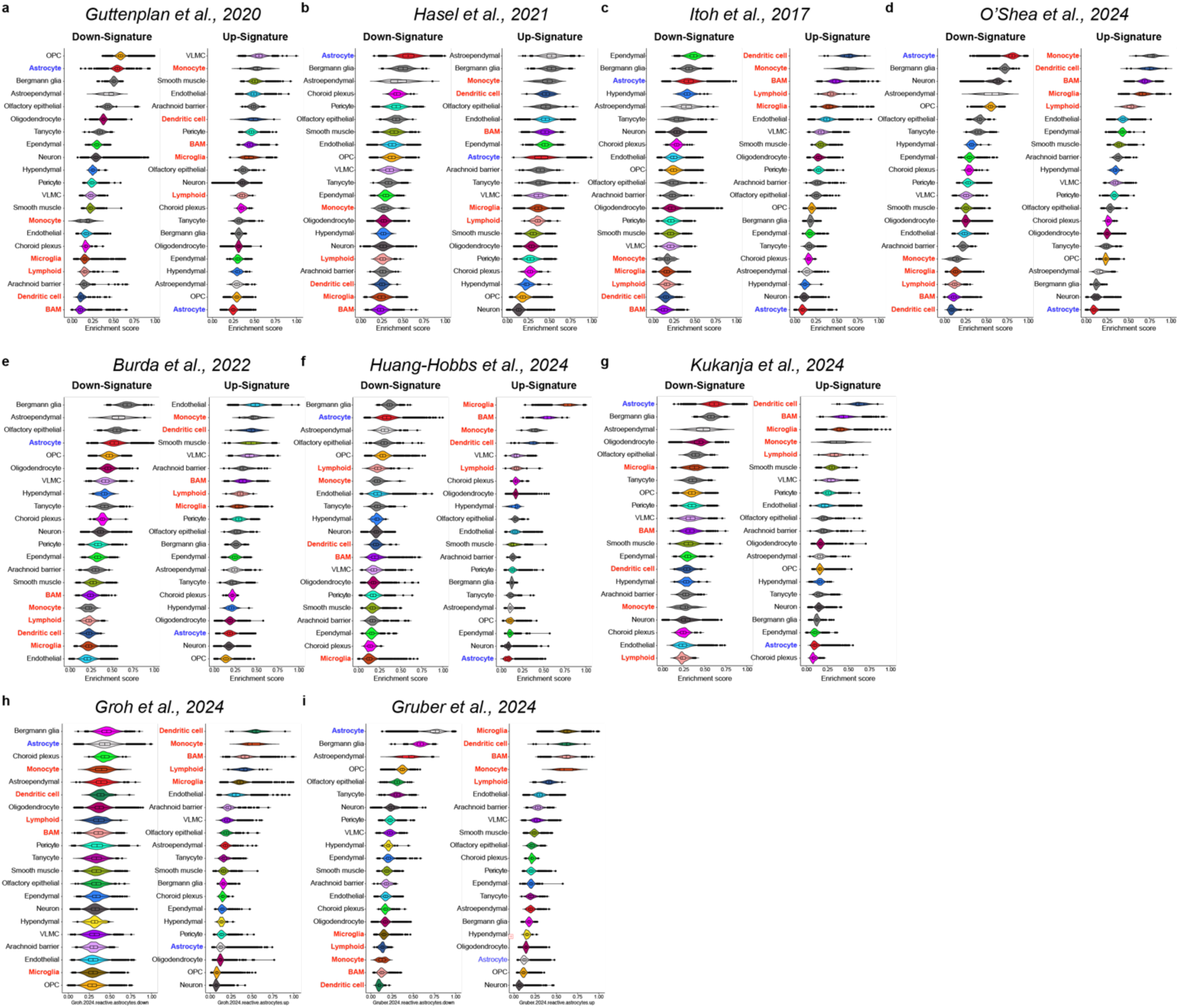
The use of Allen Brain Cell Atlas modules to study astrocytes in inflammation is misleading. **(a)** Up- and down-transcriptional signatures in immunopanned astrocytes stimulated by IL-1α+TNFα+C1q versus control, analyzed using the Allen Brain Cell Atlas cell type clusters^38^. **(b)** Up- and down-signatures in FACS-sorted astrocytes stimulated with LPS versus saline, analyzed using the Allen Brain Cell Atlas cell type clusters^33^. **(c)** Up- and down-signatures in astrocytes (EAE versus control), analyzed using the Allen Brain Cell Atlas cell type clusters^39^. **(d)** Up- and down-signatures in astrocytes (d14 SCI astrocyte versus control), analyzed using the Allen Brain Cell Atlas cell type clusters*^31^*. **(e)** Up- and down-signatures in astrocytes (LPS versus saline), analyzed using the Allen Brain Cell Atlas cell type clusters*^36^*. **(f)** Up- and down-signatures in astrocytes in glioma-bearing mice (stimulated by CNO versus Saline), analyzed usingthe Allen Brain Cell Atlas cell type clusters*^41^*. **(g)** Up- and down-signatures in astrocytes (EAE versus control), analyzed using the Allen Brain Cell Atlas cell type clusters^46^. **(h)** Up- and down-signatures in astrocytes (EAE versus control), analyzed using the Allen Brain Cell Atlas cell type clusters^47^. **(i)** Up- and down-signatures astrocytes (EAE versus control), analyzed using the Allen Brain Cell Atlas cell type clusters^48^.

**Extended Data Figure 3.**
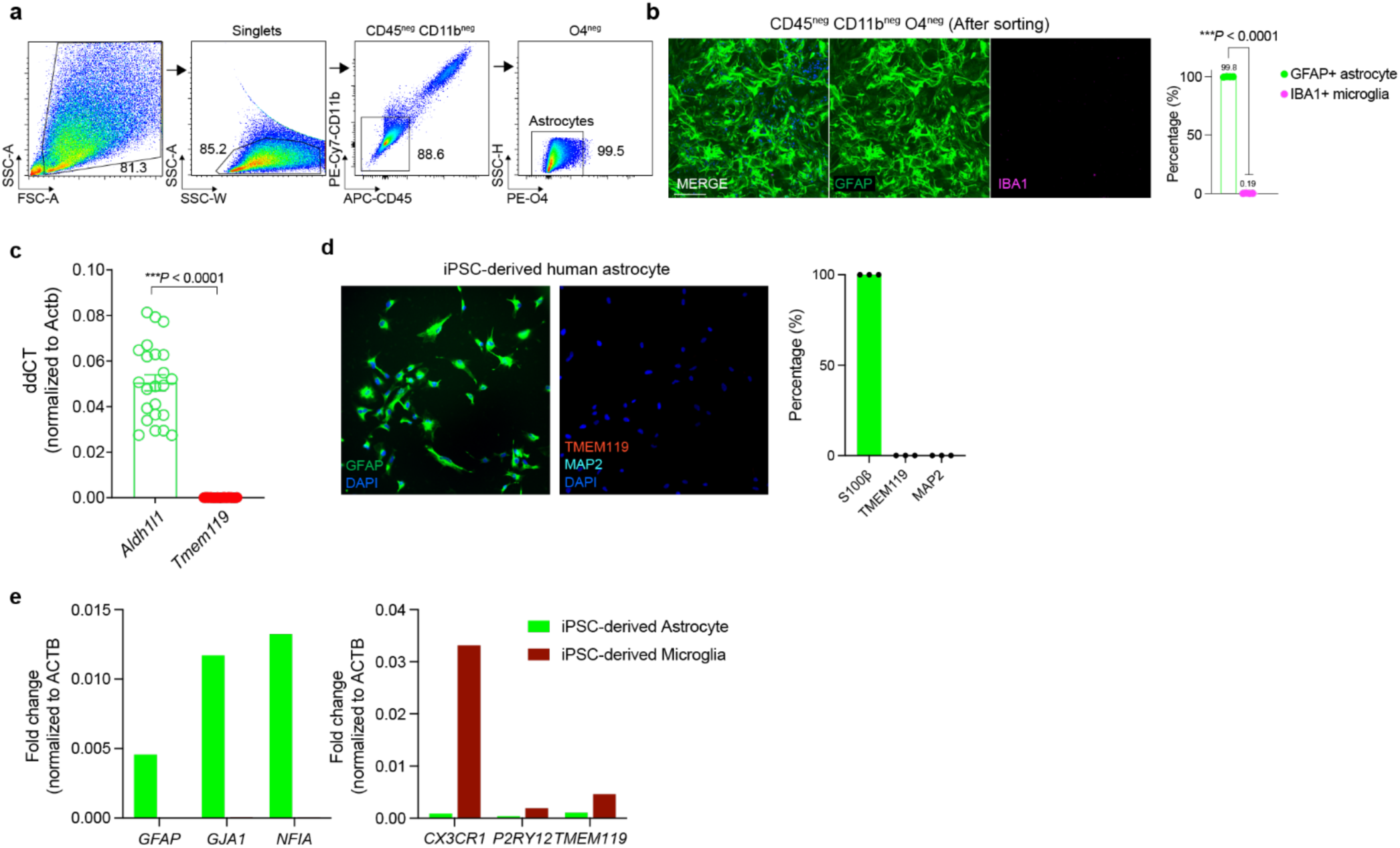
Astrocyte purity validation. **(a-b)** Purity analysis after astrocyte FACS purification based on excluding immune cells (CD45, CD11b) and oligodendrocytes (O4). Unpaired two-tailed t-test. **(c)** qPCR analysis of immunopanned astrocytes. Unpaired two-tailed t-test. **(d-e)** Purity analysis of human iPSC-derived astrocytes by immunofluorescence (d) and qPCR (e).

**Extended Data Figure 4.**
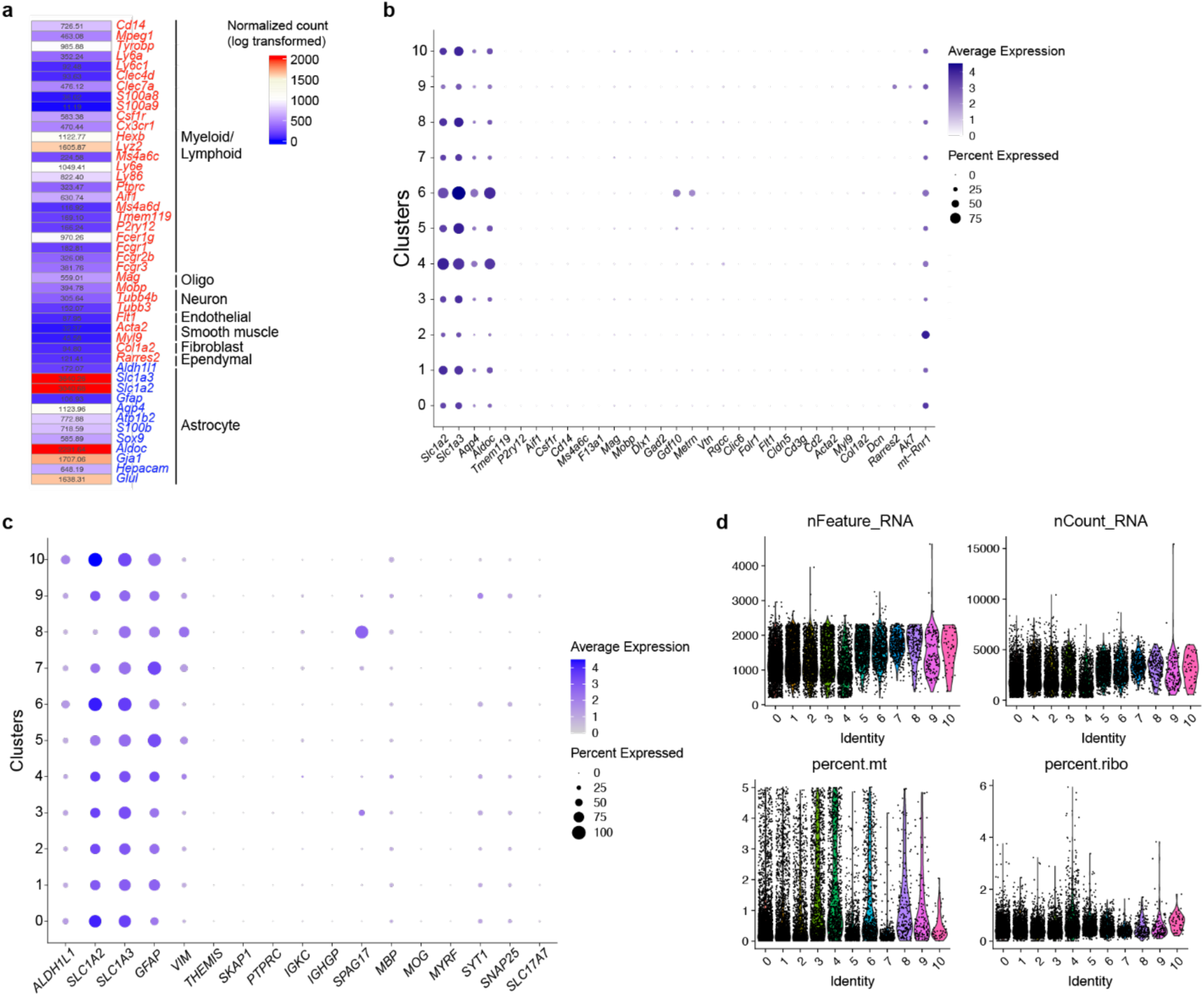
EAE/MS scRNA-seq data filtered using Liddelow lab criteria. **(a)** Pseudobulk analysis of TdTomato^Gfap^ astrocytes^105^. **(b)** TdTomato^Gfap^ astrocyte dot plot of canonical cell type marker genes across the clusters applying Hasel *et al.^33^* filtering criteria. **(c)** Astrocyte dot plot of canonical cell type marker genes across the clusters from patients with multiple sclerosis and control individuals from Schirmer *et al.*^109^ and Absinta *et al.*^110^ applying Hasel *et al.^33^* filtering criteria. **(d)** Violin plots depicting the distribution of UMI counts, detected gene numbers, and the percentage of mitochondrial and ribosomal gene expression across clusters.

**Extended Data Figure 5.**
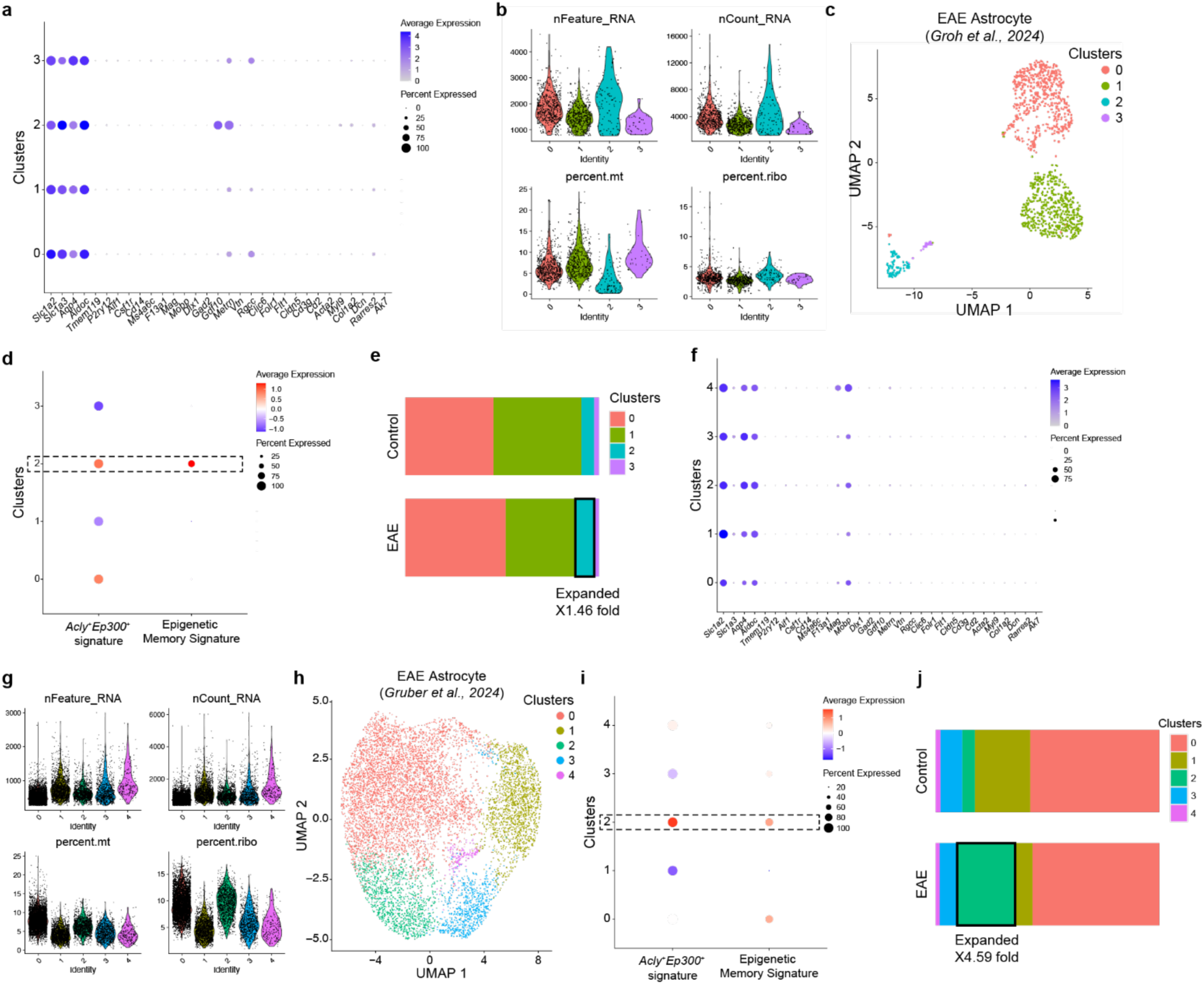
Identification of astrocyte epigenetic memory in two additional independent EAE datasets filtered using Liddelow lab criteria. **(a)** Astrocyte dot plot from Groh *et al.*^47^ of canonical cell type marker genes across clusters applying Hasel *et al.*^33^ filtering criteria. **(b)** Violin plots depicting the distribution of UMI counts, detected gene numbers, and the percentage of mitochondrial and ribosomal gene expression across clusters. **(c)** UMAP plot of astrocytes from control and EAE mice from Groh *et al.*^47^ applying Hasel *et al.*^33^ filtering criteria. **(d)** Acly^+^Ep300^+^ EAE astrocyte signature and astrocyte epigenetic memory signature expression in EAE astrocyte clusters. **(e)** Fraction of cells per cluster overall. **(f)** Astrocyte dot plot from Gruber *et al.*^48^ of canonical cell type marker genes across the clusters applying Hasel *et al.*^33^ filtering criteria. **(g)** Violin plots depicting the distribution of UMI counts, detected gene numbers, and the percentage of mitochondrial and ribosomal gene expression across clusters. **(h)** UMAP plot of astrocytes f from control and EAE mice from Gruber *et al.*^48^ applying Hasel *et al.*^33^ filtering criteria. **(i)** Acly^+^Ep300^+^ EAE astrocyte signature and astrocyte epigenetic memory signature expression in MS astrocyte clusters. **(j)** Fraction of cells per cluster overall.

**Extended Data Figure 6.**
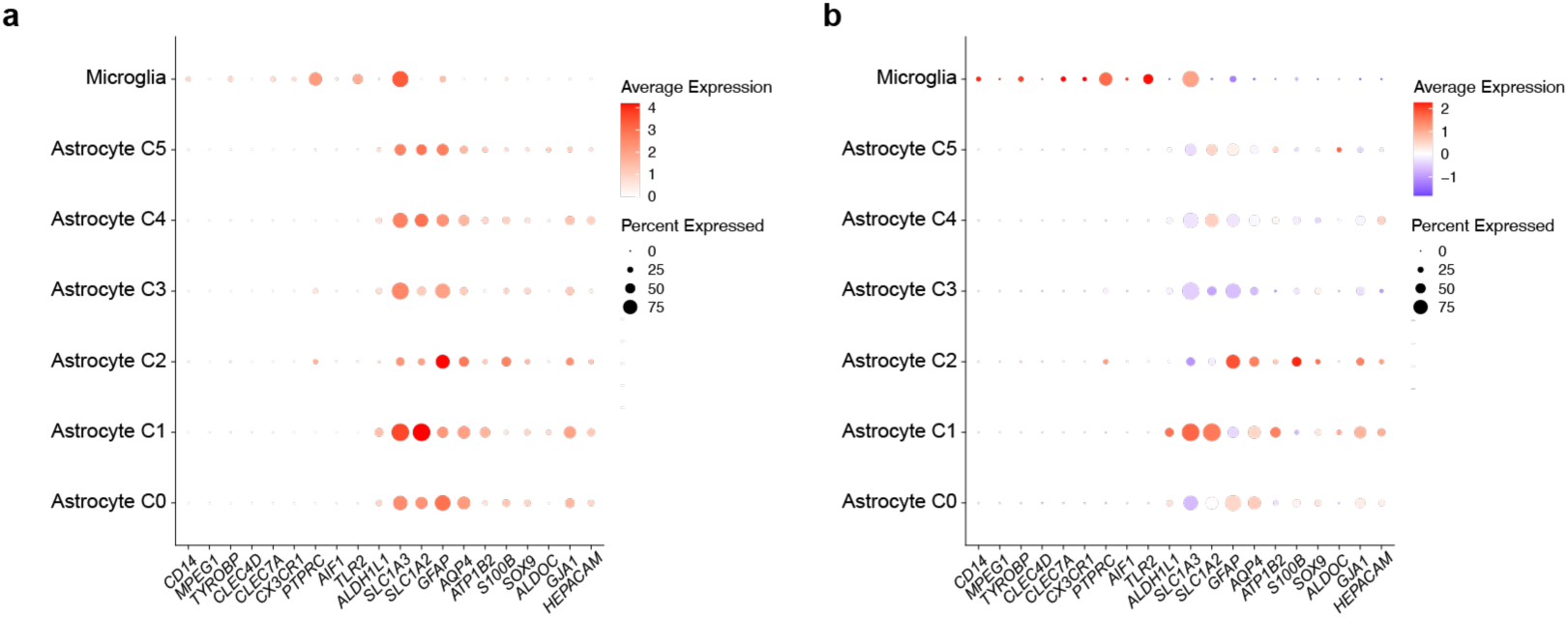
Schirmer *et al.* and Absinta *et al.* MS dataset used in Lee *et al.* **(a)** Astrocyte and microglia dot plot of canonical cell type marker genes across the clusters from patients with multiple sclerosis and control individuals from Schirmer *et al.^109^* and Absinta *et al.^110^* based on normalized counts. **(b)** Astrocyte and microglia dot plot of canonical cell type marker genes across the clusters from patients with multiple sclerosis and control individuals from Schirmer *et al.^109^* and Absinta *et al.^110^* based on scaled counts (subtracted by mean and divided by standard deviation).

**Extended Data Figure 7.**
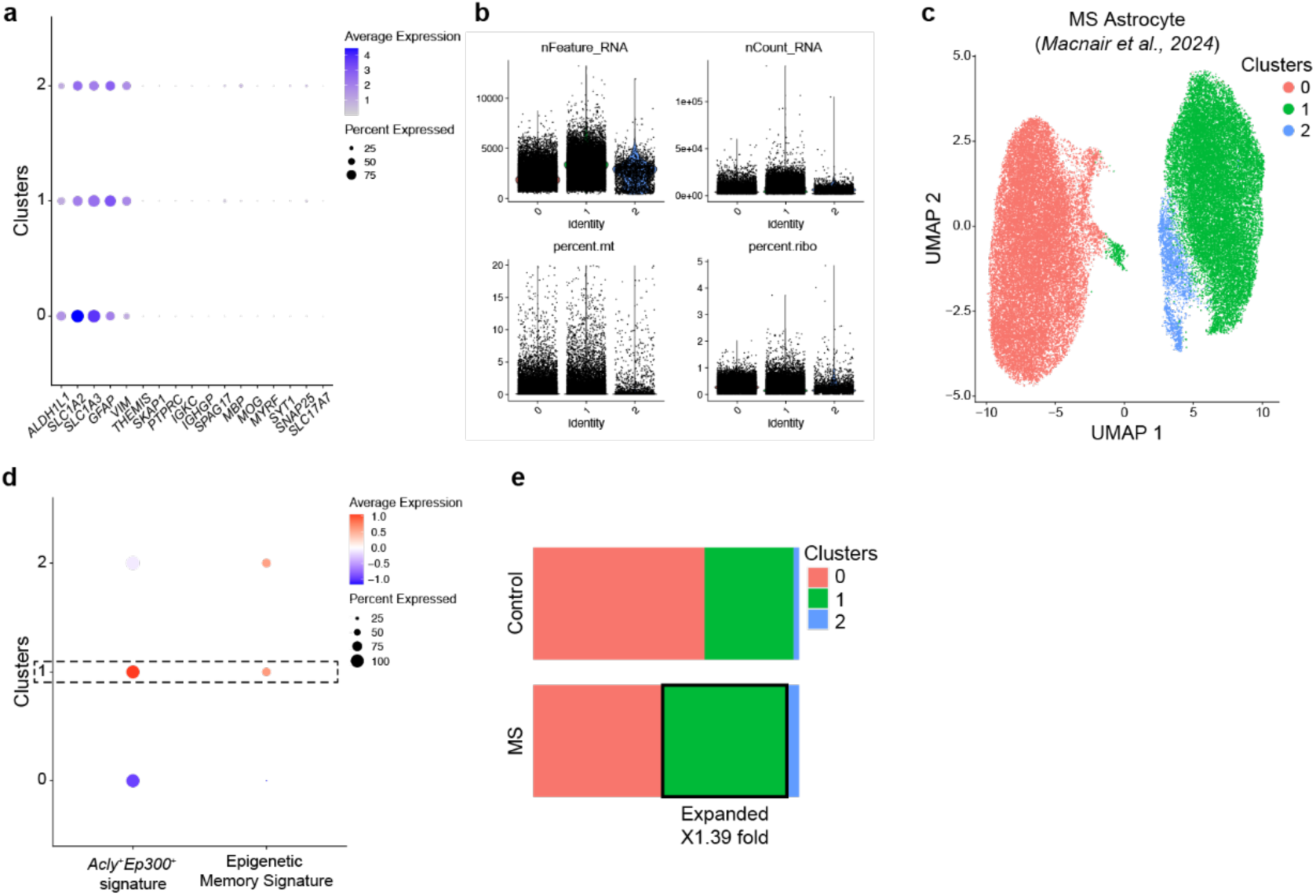
Identification of astrocyte epigenetic memory in an additional independent MS dataset filtered using Liddelow lab criteria. **(a)** Astrocyte dot plot from Macnair *et al.*^111^ of canonical cell type marker genes across the clusters applying Hasel *et al.*^33^ filtering criteria. **(b)** Violin plots depicting the distribution of UMI counts, detected gene numbers, and the percentage of mitochondrial and ribosomal gene expression across clusters. **(c)** UMAP plot of astrocytes from patients with multiple sclerosis and control individuals from Macnair *et al.*^111^ applying Hasel *et al.*^33^ filtering criteria. **(d)** Acly^+^Ep300^+^ EAE astrocyte signature and astrocyte epigenetic memory signature expression in MS astrocyte clusters. **(e)** Fraction of cells per cluster overall.

## Methods

### *In vivo* mouse astrocyte treatment and FACS purification

C57BL/6J mice at age 10-12 weeks were anesthetized using 1-3% isoflurane mixed with oxygen. Heads were shaved and cleaned using 70% ethanol and Betadine (Thermo Fisher, #19-027132) followed by a medial incision of the skin to expose the skull. Mice were injected with 100 ng/mL IL-1β (R&D Systems, #401-ML-005, 100 µg/mL stock in PBS) and 50 ng/mL TNF (R&D Systems, #410-MT-010, 100 µg/mL stock in PBS) diluted in PBS or PBS. At day 7, mice were re-injected (2X) or injected (1X) with 100 ng/mL IL-1β and 50 ng/mL TNF or PBS for 18-24 hours. The lateral ventricles were targeted bilaterally using the coordinates: +/- 1.0 (lateral), - 0.44 (posterior), -2.2 (ventral) relative to Bregma. Mice were injected by two 5µL injections using a 25 µL Hamilton syringe (Sigma-Aldrich, #20787) on a stereotaxic alignment system (Kopf, #1900) and the incision was sutured. Astrocytes were isolated by flow cytometry as described^105,122-126^ and by modifying a previously described protocol^127^. Briefly, mice were perfused with 1X PBS and the CNS was isolated into 10 mL of enzyme digestion solution consisting of 75 µL Papain suspension (Worthington, #LS003126) diluted in enzyme stock solution (ESS) and equilibrated to 37C. ESS consisted of 10 mL 10X EBSS (Sigma-Aldrich, #E7510), 2.4 mL 30% D(+)-Glucose (Sigma-Aldrich, #G8769), 5.2 mL 1M NaHCO3 (VWR, #AAJ62495-AP), 200 µL 500 mM EDTA (Thermo Fisher Scientific, #15575020), and 168.2 mL ddH2O, filter-sterilized through a 0.22 µm filter. Samples were shaken at 80rpm for 30-40 minutes at 37°C. Enzymatic digestion was stopped with 1 mL of 10X hi ovomucoid inhibitor solution and 20 µL 0.4% DNase (Worthington, #LS002007) diluted in 10 mL inhibitor stock solution (ISS). 10X hi ovomucoid inhibitor stock solution contained 300 mg BSA (Sigma-Aldrich, #A8806), 300 mg ovomucoid trypsin inhibitor (Worthington, #LS003086) diluted in 10 mL 1X PBS and filter sterilized using at 0.22 µm filter. ISS contained 50 mL 10X EBSS (Sigma-Aldrich, #E7510), 6 mL 30% D(+)-Glucose (Sigma-Aldrich, #G8769), 13 mL 1M NaHCO3 (VWR, #AAJ62495-AP) diluted in 170.4 mL ddH2O and filter-sterilized through a 0.22 µm filter. Tissue was mechanically dissociated using a 5 mL serological pipette and filtered through at 70 µm cell strainer (Fisher Scientific, #22363548) into a fresh 50 mL conical. Tissue was centrifuged at 500g for 5 minutes and resuspended in 10 mL of 30% Percoll solution (9 mL Percoll (GE Healthcare Biosciences, #17-5445-01), 3 mL 10X PBS, 18 mL ddH2O). Percoll suspension was centrifuged at 500g for 25 minutes with no brakes. Supernatant was discarded and the cell pellet was washed 1X with 1X PBS, centrifuged at 500g for 5 minutes and prepared for downstream applications. Cells were stained in the dark on ice for 20 minutes with FACS antibodies. Cells were then washed once with 1X PBS and resuspended in 1X PBS for sorting as described^105,122-126^. Antibodies used in this study were: PE anti-mouse CD45R/B220 (BD Biosciences, #553089, 1:100), PE anti-mouse TER-119 (Biolegend, #116207, 1:100), PE anti-O4 (R&D Systems, #FAB1326P, 1:100), PE anti-CD105 (eBioscience, #12-1051-82, 1:100), PE anti-CD140a (eBioscience, #12-1401-81, 1:100), PE anti-Ly-6G (Biolegend, #127608, 1:100), BV786-Ly6C (BD Biosciences, #740850, 1:100), APC anti-CD45 (eBioscience, #17-0451-83, 1:100), APC-Cy7 anti-CD11c (BD Biosciences, #561241, 1:100), and FITC anti-CD11b (eBioscience, #11-0112-85, 1:100). Astrocytes were sorted on the following parameters: CD105^neg^CD140a^neg^O4^neg^Ter119^neg^Ly-6G^neg^CD45R^neg^CD11b^neg^CD45^neg^Ly-6C^neg^CD11c^neg^.

Compensation was performed on single-stained samples of cells and an unstained control. Cells were sorted on a FACS Aria IIu (BD Biosciences) and analyzed by BD FACSDIVA (v.8.0.1). For intracellular staining cells were fixed according to the manufacturer’s protocol (eBiosciences, #00-5523-00). Intracellular antibodies used in this study were: BV421 anti-mouse GFAP (BioLegend, #644710, 1:100). FACS was performed on a Symphony A5 (BD Biosciences) and analyzed by Flowjo.

### *In vitro* primary mouse astrocyte FACS purification and treatment

Procedures were performed as described^128^. Brains of mice aged P0-P3 were dissected into PBS on ice. Cortices were discarded and the brain parenchyma were pooled, centrifuged at 500g for 10 minutes at 4°C and resuspended in 0.25% Trypsin-EDTA (Thermo Fisher Scientific, #25200-072) at 37°C for 10 minutes. Trypsin was neutralized by adding DMEM (Thermo Fisher Scientific, # 11965118) supplemented with 10% FBS (Thermo Fisher Scientific, #10438026) and 1% penicillin/streptomycin (Thermo Fisher Scientific, #15140148), and cells were passed through a 70µm cell strainer. Cells were centrifuged at 500g for 10 minutes at 4°C, resuspended in DMEM with 10% FBS/1% penicillin/streptomycin and cultured in T-75 flasks (Falcon, #353136) at 37°C in a humidified incubator with 5% CO2, for 7-10 days until confluency was reached. Astrocytes were shaken for 30 minutes at 180 rpm; the supernatant was collected to deplete microglia. Fresh medium was added, and astrocytes were shaken again at 220rpm for at least 2 hours and the supernatant was aspirated to further deplete contaminating cells. 0.25% Trypsin-EDTA was treated for 5min at 37’C and 5% CO2 to detach the astrocytes. Trypsin was neutralized by adding DMEM/F12+GlutaMAX supplemented with 10% FBS and 1% penicillin/streptomycin, and astrocytes were pelleted and washed. Washed astrocyte pellet was treated with APC anti-CD45 (Biolegend, #103112), PECy7 anti-CD11b (Biolegend, #101215) and PE anti-Olig4 (R&D systems, # FAB1326P). Astrocytes were further purified by sorting CD45-CD11b-Olig4-population and plated on poly-L-lysine (Sigma, #P4707) pre-coated plates. Pre-sorted (CD45-CD11b-Olig4-) astrocytes were treated with 100 ng/mL IL-1β (R&D Systems, #401-ML-005, 100 µg/mL stock in PBS) and 50 ng/mL TNF (R&D Systems, #410-MT-010, 100 µg/mL stock in PBS) diluted in DMEM (Thermo Fisher Scientific, # 11965118) that was supplemented with 10% FBS (Life Technologies, #10438026) and 1% penicillin/streptomycin (Life Technologies, #15140122) or PBS. The medium was replaced and washed twice after 18-24 hours, then astrocytes were rested for 6 days. Medium was replaced every 2–3 days. At day 7, astrocytes were re-treated (2X) or treated (1X) with 100 ng/mL IL-1β and 50 ng/mL TNF or PBS for 30min. For p65 sorting experiments, EGFP^pos^ and EGFP^neg^ astrocytes were FACS sorted after 18-24 hours of IL-1β and TNF treatment and re-seeded and rested for 6 days. Astrocytes were subsequently gated on: CD11b^neg^CD45^neg^EGFP^pos/neg^. Medium was replaced every 2–3 days. At day 7, astrocytes were re-treated (2X) or treated (1X) with 100 ng/mL IL-1β and 50 ng/mL TNF or PBS for 30min.

### *In vitro* primary mouse astrocyte immunopanning and treatment

Astrocyte isolation by immunopanning was performed following the protocol developed by Foo et al*^129,130^*. Briefly, mouse pups aged <p7 were cryoanesthetized and euthanized by decapitation. Cortices were dissected out, minced and enzymatically dissociated with 100µl Papain (Worthington Biochemical, LS003126), 200µl DNase (Thermo Fisher Scientific, 90083) and 4mg L-Cysteine (Sigma-Aldrich, C7880) in 20ml enzyme stock solution (ESS) made with 20ml 10X EBSS (Sigma-Aldrich, E7510), 2.4ml 30% D-Glucose (Sigma-Aldrich, G8769), 5.2ml 1M NaHCO3 (VWR, AAJ62495-AP), 200µl 0.5M EDTA (Thermo Fisher Scientific, 15575020) and 172.2ml ddH2O (Thermo Fisher Scientific, 10-977-023) for 45 min under 37°C and 5% CO2. Brains chunks were transferred to a tube and the supernatant carefully removed. Enzymatic digestion was then stopped by adding 5ml “Low” Ovomucoid Trypsin Inhibitor solution in inhibitor stock solution (ISS). To prepare this solution, a 10X Ovomucoid inhibitor stock was prepared with 0.15g Ovomucoid inhibitor (Worthington Biochemical, LS003087), 0.15g BSA (Sigma-Aldrich, A3294) and 10ml DPBS (Thermo Fisher Scientific, 14-190-250). ISS was made with 50ml 10X EBSS, 6ml 30% D-Glucose, 13ml NaHCO3 and 431ml ddH2O. After letting brain chunks settle, supernatant was removed and new “Low” Trypsin inhibitor solution was added, repeating the process four times. Finally, brain tissue was triturated by pipetting several times. A “High” Trypsin inhibitor solution was prepared from a 10X concentrated stock made with 0.3g Ovomucoid inhibitor, 0.3g BSA and 10ml DPBS, and dissolved in ISS. 10ml “High” Trypsin inhibitor solution was pipetted below the cell suspension in order to create a gradient. This suspension was centrifuged for 5 min at 800 rpm, resuspended in 10ml 0.2% BSA in PBS, and filtered through 70µm strainer (Celltreat, 229484), washing through the filter with an additional 10ml 0.2% BSA. Cell suspensions were then rested for 45 mins at 37°C in a water bath prior to immunopanning. Panning plates were coated overnight with secondary antibodies in Tris-HCl pH 9.5 (Boston BioProducts, BBT-95), using 60µl polyclonal Goat anti-Rat IgG antibody (Jackson ImmunoResearch, 112-005-167) or 60µl goat anti-Mouse IgG+IgM (Jackson ImmunoResearch, 115-005-044), respectively. Additional plates were coated with 50µg per plate Griffonia (Bandeiraea) Simplicifolia Lectin I (BSL-1) (Vector Laboratories, L-1100-5). Several hours before immunopanning, coated plates were washed with DPBS three times, and primary antibodies diluted in 0.2% BSA-DPBS were added. Primary antibodies were mouse IgM anti-O4 (R&D Systems, MAB1326), rat IgG anti-CD45 (BD Pharmingen, 550539), mouse IgG1 anti-HepaCAM (R&D Systems, MAB4108). Prior to immunopanning, the coated plates were washed with DPBS three times, while saving the anti-HepaCAM solution for later use. Cell suspensions were sequentially added to the anti-O4 plate for 20 mins at room temperature (RT), then onto the second anti-O4 plate for 20 mins at RT, then onto the anti-CD45 plate for 20 mins at RT, and then onto the unwashed BSL-1 plate for 10 minutes. Finally, the remaining cell suspension was mixed with the anti-HepaCAM solution to maximize binding of astrocytes and incubated at 37°C in a shaker for 30 mins. Meanwhile, the anti-HepaCAM plate was washed three times with DPBS. Cell suspensions were then added to the anti-HepaCAM plate for 30 mins at RT. Next, the positive selection plate was gently washed with DPBS until no floating cells were visible under the microscope. To detach cells, 200U of trypsin (Sigma-Aldrich, T9935) in 8ml of 1X EBSS were added and plates lightly tapped. To stop dissociation and collect all cells, 10ml of 30% fetal bovine serum (FBS) (Thermo Fisher Scientific, A5670801) in Neurobasal A medium (Thermo Fisher Scientific, 12349015) were pipetted all around the plate. This was repeated with another 10ml 30% FBS. Cell suspensions were then diluted with an equal amount of 0.02% BSA-DPBS and centrifuged at 1000rpm for 11 mins. Culture plates were coated with poly-D-lysine (Thermo Fisher Scientific, A3890401) for 1h and washed with DPBS. Cells were counted using a hemacytometer and plated in astrocyte culture medium prepared with 144ml Neurobasal A, 144ml DMEM (Thermo Fisher Scientific, 11965118), 3ml Penicillin/Streptomycin (Thermo Fisher Scientific, 15140122), 3ml 100mM Sodium Pyruvate (Gibco, 11360070), 3ml 100X GlutaMAX (Thermo Scientific, 35050061), 3 ml 100X SATO, and 300µl 5mg/ml N-Acetyl-cysteine (Sigma-Aldrich, A7250). 100X SATO was prepared with 6ml Neurobasal A, 60mg transferrin (Sigma-Aldrich, T8158), 60mg BSA, 9.6mg putrescine (Sigma-Aldrich, P5780), 1.5µl 25mg/ml progesterone (Sigma-Aldrich, P8783), 60µl 0.4mg/ml sodium selenite (Sigma-Aldrich, S5261). Astrocyte cultures were supplemented with recombinant HB-EGF (Thermo Fisher Scientific, 100-47) to 5ng/ml final concentration. Immunopanned astrocytes were treated with 100 ng/mL IL-1β (R&D Systems, #401-ML-005, 100 µg/mL stock in PBS) and 50 ng/mL TNF (R&D Systems, #410-MT-010, 100 µg/mL stock in PBS) or PBS. The medium was replaced and washed twice after 18-24 hours, then astrocytes were rested for 6 days. Medium was replaced every 2–3 days. At day 7, astrocytes were re-treated (2X) or treated (1X) with 100 ng/mL IL-1β and 50 ng/mL TNF or PBS for 1h. p300 inhibition was performed with 1 μM C646 (SelleckChemicals S7152)for 2h at 37 °C before and during the first cytokine treatment

### iPSC-derived astrocyte differentiation and treatment

iPSC astrocyte cell line ND48, a gift from Vikram Khurana, was used for all iPSC astrocyte experiments. ND48 was differentiated following a modified protocol^131^. On day 0, IPSCs at 80-90% confluency was dissociated with accutase (Life Technologies, 00-4555-56), and 6 x 10^5^ cells/cm^2^ were plated in Essential 6 medium (Thermo Fisher Scientific, A1516401) with 10 μM ROCK inhibitor (StemCell Technologies, Y-27632). On days 1-2, cells were fed with Essential 6 medium supplemented with 2 μM XAV 939 (Tocris Bioscience, 3748), 100 nM LDN (Tocris Bioscience, 6053), and 10 μM SB (Tocris Bioscience, 1614). On days 3-9, cells were fed with Essential 6 medium supplemented with 100nM LDN and 10 μM SB. On days 10-16, cells were fed with medium containing 50% Neurobasal medium (Thermo Fisher Scientific, 21103049), 50% DMEM/F12 medium (Thermo Fisher Scientific, 11320033), 2% B27 supplement (Thermo Fisher Scientific, 17504044), 1% N2 supplement (Thermo Fisher Scientific, 17502048), and 1% penicillin-streptomycin (Thermo Fisher Scientific, 15070063) (N2/B27 medium). On day 16, cells were dissociated with accutase, and 2 x 10^5^ cells/cm^2^ were plated on Matrigel-coated 6-well plates in N2/B27 medium containing 10 μM ROCK inhibitor. On day 17, cells were fed with N2/B27 medium. On day 18, cells were transduced with 2 μg FUW-NFIA and 2 μg FUW M2-rtTA lentiviruses in N2/B27 medium. Lentivirus packaging was done by Boston Children’s Hospital Viral Core Facility. On days 19-24, cells were fed with N2/B27 medium containing 1 µg/mL doxycycline hyclate (Stemcell Technologies, 72742). On days 25-50, cells were propagated with daily media changes of astrocyte medium (ScienCell, 1801) with 2% FBS and passaged at 80-90% confluency (complete AM). On day 3, iPSC-derived astrocytes were plated on Matrigel coated 48 well plates at 1 x 10^4^ cell/ well in complete AM. On day 0, the cells were treated for 24 h with 10 ng ml^−1^ IL-1β (Gibco, PHC0815) and 5 ng ml^−1^ TNF (Gibco, PHC3016) in complete AM. After the first stimulation, cells were washed 2 times with warmed complete AM and then cultured for the subsequent 6 days. On day 7, iPSC-derived astrocytes were re-treated (2X) or treated (1X) with 10 ng/ml IL-1β and 5 ng/ml TNF or PBS for 2 h. For the control conditions, the media aspiration, media change and washing occurred alongside all treated cells.

### RNA expression analysis of primary astrocytes

Primary astrocytes were lysed in Buffer RLT (Qiagen) and RNA was isolated from cultured astrocytes using the Qiagen RNeasy Mini kit (Qiagen, #74106). cDNA was transcribed using the High-Capacity cDNA Reverse Transcription Kit (Life Technologies, #4368813). Gene expression was then measured by qPCR using Taqman Fast Universal PCR Master Mix (Life Technologies, #4367846). Taqman probes used in this study are: *Actb* (Mm02619580_g1), *Nos2* (Mm00440502_m1), *Ccl2* (Mm00441242_m1), *Il6* (Mm00446190_m1), *Aldh1l1* (Mm03048957_m1), *Tmem119* (Mm00525305_m1), *ACTB* (Hs01060665_g1), *IL6* (Hs00174131_m1), *GFAP* (Hs00909233_m1), *GJA1* (Hs00748445_s1), *NFIA* (Hs00325656_m1), *TMEM119* (Hs01938722_u1), *CX3CR1* (Hs01922583_s1), *P2RY12* (Hs01881698_s1), *IL1B* (Hs01555410_m1), and *TNF* (Hs00174128_m1). qPCR data were analyzed by the ddCt method by normalizing the expression of each gene for each replicate to β-actin and then to the control group.

### Immunostaining of mouse primary astrocytes

Fluorescence-activated cell sorted astrocytes were plated on poly-L-lysine coated optical bottom plates (Thermo Scientific, #165305), and incubated for 2-3days. Astrocytes were washed 2 times with 1x PBS (Gibco, #14-190-250) and fixed with 4% PFA (EMS, #15714) in 1x PBS for 10min at room temperature. Astrocytes were washed 2 times with 1x PBS, and blocked with 4% bovine serum albumin (Sigma-Aldrich, A3294) in 0.3% Triton X-100 at room temperature for an hour. Astrocytes were then incubated with primary antibody (rat anti-GFAP, Invitrogen, #13-0300 & rabbit anti-IBA1, Wako, #019-19741) diluted in blocking buffer overnight at 4 °C. Following primary antibody incubation, sections were washed 3 times with 0.1% PBS-T and incubated with secondary antibody (Alexa Flour 647 conjugated goat anti-rat IgG, Thermo Fisher Scientific, #A21247 & Alexa Flour 555 conjugated donkey anti-rabbit IgG, Thermo Fisher Scientific, #A31572) diluted in 0.1% PBS-T for 2 h at room temperature. Sections were then washed 3 times with 0.1% PBS-T and mounting media with DAPI (Thermo Scientific, #P36931) was dropped on stained tissue. Images were obtained by LSM880 (Zeiss).

### Immunostaining of human primary astrocytes

Human astrocytes were plated on poly-L-lysine coated optical bottom plates (Thermo Scientific, #165305), and incubated for 2-3days. Astrocytes were washed 2 times with 1x PBS (Gibco, #14-190-250) and fixed with 4% PFA (EMS, #15714) in 1x PBS for 10min at room temperature. Astrocytes were washed 2 times with 1x PBS, and blocked with 4% bovine serum albumin (Sigma-Aldrich, A3294) in 0.3% Triton X-100 at room temperature for an hour. Astrocytes were then incubated with primary antibody (rat anti-GFAP, Invitrogen, #13-0300 & rabbit anti-TMEM119, Millipore Sigma, #HPA051870) diluted in blocking buffer overnight at 4 °C. Following primary antibody incubation, sections were washed 3 times with 0.1% PBS-T and incubated with secondary antibody (Alexa Flour 647 conjugated goat anti-rat IgG, Thermo Fisher Scientific, #A21247 & Alexa Flour 555 conjugated donkey anti-rabbit IgG, Thermo Fisher Scientific, #A31572) diluted in 0.1% PBS-T for 2 h at room temperature. Sections were then washed 3 times with 0.1% PBS-T and mounting media with DAPI (Thermo Scientific, #P36931) was dropped on stained tissue. Images were obtained by LSM880 (Zeiss).

### Immunostaining of human iPSC-derived astrocytes

Immunofluorescence analysis was performed as follows. IPSC-derived astrocyte cultures grown in 96-well clear bottom plates (Thermo Fisher Scientific, 165305) were fixed with 100 μL of 4% paraformaldehyde in PBS for 15 min. Cells were washed three times with PBS, 5 min per wash. Cells were blocked and permeabilized using 10% goat serum (Thermo Fisher Scientific, 16210064), 0.1% Saponin (to preserve integrity of lipid-rich inclusions) (Boston Bioproducts, BM-688) in PBS for 1 h at room temperature. Primary antibody was incubated in 2% goat serum, 0.02% Saponin overnight at 4°C. Cells were washed three times with PBS, 5 min per wash, and incubated with secondary antibody in 2% goat serum, 0.02% Saponin and 1% DAPI for 1 h at 37°C. Finally, cells were washed three times with PBS, 5 min per wash. Images of the immunostained cells were captured with a Nikon TiE/C2 confocal microscope. Primary antibodies were mouse anti-S100β (MilliporeSigma, S2532, 1:500), chicken anti-MAP2 (Abcam, ab92434, 1:1000), mouse anti-TMEM119 (BioLegend, 853302, 1:1000). Secondary antibodies were: donkey anti-mouse IgG (H+L), Alexa Fluor 488 (Thermo Fisher Scientific, A21202), donkey anti-chicken IgY (H+L), Alexa Fluor 488 (Thermo Fisher Scientific, A78948), donkey anti-mouse IgG (H+L), Alexa Fluor 594 all at 1:500.

### Heatmap and Allen Brain Cell Atlas analysis of Liddelow *et al.*, and other datasets

The bulk RNA-seq of astrocytes samples from the following publications were analyzed: Guttenplan *et al.*^38^, Itoh *et al.^39^,* Burda *et al.*^36^, O’Shea *et al.*^31^, and Sardar *et al.*^40^. The raw count matrix downloaded from each GEO repository was normalized by the counts-per-million approach implemented by edgeR^132^, and then log transformed. The log-transformed normalized count matrices were visualized using ComplexHeatmap^133^. Differential expression analyses were performed using DESeq2^134^ between reactive and control condition, genes that have log2-fold change larger than 1 and adjusted p-value (the DESeq2 uses false discovery rate to generate the adjusted p-value to counter issues brought by multiple testing) smaller than 0.05 are deemed as “up-regulated genes in reactive astrocytes” and genes that have log2-fold change smaller than (-1) and adjusted p-value smaller than 0.05 are deemed as “down-regulated genes in reactive astrocytes”. In Guttenplan *et al.*^38^, the differential expression results were uploaded, so the genes were directly selected from the given results. Spatial transcriptomic and single-cell RNA-seq datasets of astrocytes samples from the following publications were analyzed: Hasel *et al.*^33^, Huang-Hobbs *et al*.^41^, Groh *et al*.^47^, Gruber *et al*.^48^, and Kukanja^46^. In Huang-Hobbs *et al*^41^, Groh *et al*.^47^, Gruber *et al*.^48^, and Kukanja *et al.*^46^ the cell type annotation was provided by the authors, so astrocytes were identified and ready to be used. In Hasel *et al.* the count matrix objects (10X format) were downloaded from the public data repository, and DoubletFinder^135^ was used to identify doublets in the dataset. Then cells with less UMIs than the 25 percentiles of UMIs or less genes than the 25 percentiles of genes detected in the sample was filtered out, and cells with more than 25% mitochondrial contents or more than 25% ribosomal contents were removed. Harmony^136^ was used for batch effect correction, and Seurat^137^ was used to do the normalization (NormalizeData function^137^), dimension reduction (RunPCA function^137^) and clustering (FindClusters function^137^). The top 15 harmony-corrected^111^ principal components were used to generate the UMAP dimension reduction (RunUMAP^112^) and build the k-nearest neighborhood results (RunNeighobors^112^); finally FindClusters^112^ was used to identified 7 clusters using resolution of 0.1 in the dataset. Clusters of cells ultimately filtered out were identified according to the criteria originally specified in Hasel *et al.*, which used canonical cell type markers to filter out clusters co-expressing markers of astrocytes and other cell types, specifically: *Flt1*/*Cldn5* as endothelial cells, *Olig1*/*Mog*/*Mobp* as oligodendrocytes, *Cd14*/*Cx3cr1*/*Tmem119*/*Lyz2* as myeloid cells, *Syt1*/*Rbfox2* as neurons. After filtering out these cells, the remainder were analyzed using canonical astrocyte markers: *Gfap*, *Gja*, *Aldoc*, *Aldh1l1*, *Slc1a2* and *Slc1a3* with negligible expression of the genes listed above. Differential expression analyses between astrocytes in stimulated sample(s) and the ones in control sample(s) in all the datasets were performed using FindMarkers^137^ function in Seurat, “up-regulated genes in reactive astrocytes” and “down-regulated genes in reactive astrocytes” were identified based on the FindMarkers results. The AggregateExpression^137^ in Seurat was used to generate pseudo-bulk of astrocytes in each sample, and ComplexHeatmap was used to visualize the expression of astrocytes and non-astrocytes markers in each pseudo-bulk sample. For the “up-regulated genes in reactive astrocytes” and “down-regulated genes in reactive astrocytes” identified in the different datasets, the single cell RNA-seq dataset of mouse brain curated by Allen Brain Cell Atlas was downloaded^42^. The dataset was in the format of a Seurat object, with comprehensive cell type annotation. The AddModuleScore^137^ function was used to calculate the module scores of the “up-regulated genes in reactive astrocytes” and “down-regulated genes in reactive astrocytes” identified in the different datasets. This function has been used and validated in multiple publications by the Regev lab and others^138-141^. The module scores were visualized by R package ggpubr.

### Reanalysis of astrocyte scRNA-seq data

Single-cell RNA-seq datasets were reanalyzed from previously published studies applying Hasel *et al.*^33^ filtering criteria., including EAE (Wheeler *et al.*^105^, Groh *et al.*^47^, and Gruber *et al.*^48^) and MS (Schirmer *et al.^109^*, Absinta *et al.^110^*, and Macnair *et al.*^111^). Among these, datasets from Wheeler *et al.*, Schirmer *et al.*, and Absinta *et al.* were previously utilized in Lee *et al.*, whereas Groh *et al.,* Gruber *et al.,* and Macnair *et al.* represent independent datasets analyzed in this study. Astrocytes previously identified by the authors of corresponding publications were sub-clustered using Seurat. The data was first normalized, and then the variable features in each sample was found using FindVariableFeatures^137^. Then SelectIntegrationFeatures^137^ was used to select the variable features used in downstream analysis for all samples. RunPCA^137^ was used to do the dimension reduction, and Harmony was used to correct the batch effect based on the results of it. Then RunUMAP^112^ was used to generate the UMAP dimension reduction, and FindNeighbors^137^ and FindClusters were used to identify the clusters. During the analysis, the top 15, 6, 13, 20 and 20 principal components were used to generate UMAP and nearest-neighborhood graphs for Macnair *et al*., Gruber *et al*., Groh *et al*., Lee *et al*., and Wheeler *et al*. single cell datasets respectively; resolutions of 0.1, 0.2, 0.6, 0.2 and 0.2 were used to do the clustering for Macnair *et al*., Gruber *et al*., Groh *et al*., Lee *et al*., and Wheeler *et al*. single cell datasets respectively. The clustering results were used to identify potential non-astrocytes contamination and low-quality cells. We filtered out cells as in the mouse analyses stated above and previously performed in Hasel *et al.*^33^ by canonical cell type markers: *TMEM119*/*P2RY12*/*AIF1*/*CSF1R*/*CD14* for myeloid cells, *CD2*/*CD3D*/*CD3E*/*CD3G* for T cells, *MOG*/*MBP* for oligodendrocytes. The number of genes and number of UMIs in each cell in each cluster were plotted out using violin plot, and clusters with significantly less genes and UMIs relative to other clusters were identified as “low quality cells”. After removing the low-quality cell population and the non-astrocytes, the rest of the true astrocytes were clustered again. The epigenetic memory signature and *Acly+Ep300*+ signature were calculated in the sub-clustered astrocytes using AddModuleScore. This function has been used and validated in multiple publications by the Regev lab and others^138-141^.

### Pathway analysis

To determine regulators of gene expression networks, Ingenuity Pathway Analysis software (Qiagen) was used by inputting gene expression datasets with corresponding log (Fold Change) expression levels compared to other groups. “Canonical pathways” and “upstream analysis” metrics were considered significant at p-value <0.05. Gene sets used for signature scores are indicated in the corresponding figure.

## Code availability

The code used to perform all bioinformatic analyses in this manuscript was deposited to Zenodo and is available under: https://zenodo.org/records/15288719?token=eyJhbGciOiJIUzUxMiIsImlhdCI6MTc0NTg1ODczMSwiZ XhwIjoxNzc3MzM0Mzk5fQ.eyJpZCI6IjEyMDg0M2IwLWI0MTktNGQxMy05ZjEzLWU5ZjkwYzJiO GUzZiIsImRhdGEiOnt9LCJyYW5kb20iOiJjZjUzOTI4NDRkOGRhNDIzZGE4OTEyNTAzNmQwMj Y2MCJ9.NftlDx4CxCx7LWPM61tEIdWtsGlGfl7W6mOjal6FrBSvdbDORlLBxegLzRckPzQQ8SAd2a N8MAyUb0ymBOL2zg.

## Authors’ contribution

H-G.L., V.R., M.A.W. and F.J.Q. designed research; H-G.L., C.F.A., J-H.L. and G.P. performed experiments; H-G.L., Z.L. and C.F.A. performed bioinformatic analyses; V.R., J.A., A.P. and I.C.C. discussed and/or interpreted findings; H-G.L., M.A.W. and F.J.Q. wrote the paper with input from coauthors.

## Competing interests

The authors declare no competing interests.

## Notes

### Competing Interest Statement

The authors have declared no competing interest.

## References

1 Allen, N. J. & Lyons, D. A. Glia as architects of central nervous system formation and function. Science 362, 181–185 (2018). 10.1126/science.aat0473

2 Lee, H. G., Wheeler, M. A. & Quintana, F. J. Function and therapeutic value of astrocytes in neurological diseases. Nat Rev Drug Discov 21, 339–358 (2022). 10.1038/s41573-022-00390-x

3 Sofroniew, M. V. Astrocyte barriers to neurotoxic inflammation. Nat Rev Neurosci 16, 249–263 (2015). 10.1038/nrn3898

4 Vainchtein, I. D. et al. Astrocyte-derived interleukin-33 promotes microglial synapse engulfment and neural circuit development. Science 359, 1269–1273 (2018). 10.1126/science.aal3589

5 Linnerbauer, M., Wheeler, M. A. & Quintana, F. J. Astrocyte Crosstalk in CNS Inflammation. Neuron 108, 608–622 (2020). 10.1016/j.neuron.2020.08.012

6 Yu, X. & Khakh, B. S. SnapShot: Astrocyte interactions. Cell 185, 220–220.e221 (2022). 10.1016/j.cell.2021.09.029

7 Rothhammer, V. et al. Type I interferons and microbial metabolites of tryptophan modulate astrocyte activity and central nervous system inflammation via the aryl hydrocarbon receptor. Nat Med 22, 586–597 (2016). 10.1038/nm.4106

8 Alaamery, M. et al. Role of sphingolipid metabolism in neurodegeneration. J Neurochem 158, 25–35 (2020). 10.1111/jnc.15044

9 Clark, I. C. et al. Barcoded viral tracing of astrocyte networks in CNS inflammation. Science 372, eabf1230 (2021).

10 Liddelow, S. A. et al. Neurotoxic reactive astrocytes are induced by activated microglia. Nature 541, 481–487 (2017). 10.1038/nature21029

11 Wheeler, M. A. et al. Environmental Control of Astrocyte Pathogenic Activities in CNS Inflammation. Cell 176, 581–596.e518 (2019). 10.1016/j.cell.2018.12.012

12 Wheeler, M. A. et al. Droplet-based forward genetic screening of astrocyte-microglia cross-talk. Science 379, 1023–1030 (2023). 10.1126/science.abq4822

13 Endo, F. et al. Molecular basis of astrocyte diversity and morphology across the CNS in health and disease. Science 378, eadc9020 (2022). 10.1126/science.adc9020

14 Zinkernagel, R. M. et al. On immunological memory. Annu Rev Immunol 14, 333–367 (1996). 10.1146/annurev.immunol.14.1.333

15 Wendeln, A. C. et al. Innate immune memory in the brain shapes neurological disease hallmarks. Nature 556, 332–338 (2018). 10.1038/s41586-018-0023-4

16 Saeed, S. et al. Epigenetic programming of monocyte-to-macrophage differentiation and trained innate immunity. Science 345, 1251086 (2014). 10.1126/science.1251086

17 Sun, J. C., Beilke, J. N. & Lanier, L. L. Adaptive immune features of natural killer cells. Nature 457, 557–561 (2009). 10.1038/nature07665

18 Serafini, N. et al. Trained ILC3 responses promote intestinal defense. Science 375, 859–863 (2022). 10.1126/science.aaz8777

19 Naik, S. et al. Inflammatory memory sensitizes skin epithelial stem cells to tissue damage. Nature 550, 475–480 (2017). 10.1038/nature24271

20 Novakovic, B. et al. beta-Glucan Reverses the Epigenetic State of LPS-Induced Immunological Tolerance. Cell 167, 1354–1368 e1314 (2016). 10.1016/j.cell.2016.09.034

21 Cheng, S. C. et al. mTOR- and HIF-1alpha-mediated aerobic glycolysis as metabolic basis for trained immunity. Science 345, 1250684 (2014). 10.1126/science.1250684

22 Netea, M. G. et al. Defining trained immunity and its role in health and disease. Nat Rev Immunol 20, 375–388 (2020). 10.1038/s41577-020-0285-6

23 Ordovas-Montanes, J. et al. Allergic inflammatory memory in human respiratory epithelial progenitor cells. Nature 560, 649–654 (2018). 10.1038/s41586-018-0449-8

24 Krausgruber, T. et al. Structural cells are key regulators of organ-specific immune responses. Nature 583, 296–302 (2020). 10.1038/s41586-020-2424-4

25 Gonzales, K. A. U. et al. Stem cells expand potency and alter tissue fitness by accumulating diverse epigenetic memories. Science 374, eabh2444 (2021). 10.1126/science.abh2444

26 Tough, D. F., Tak, P. P., Tarakhovsky, A. & Prinjha, R. K. Epigenetic drug discovery: breaking through the immune barrier. Nat Rev Drug Discov 15, 835–853 (2016). 10.1038/nrd.2016.185

27 Larsen, S. B. et al. Establishment, maintenance, and recall of inflammatory memory. Cell Stem Cell 28, 1758–1774.e1758 (2021). 10.1016/j.stem.2021.07.001

28 Lee, H. G. et al. Disease-associated astrocyte epigenetic memory promotes CNS pathology. Nature 627, 865–872 (2024). 10.1038/s41586-024-07187-5

29 O’Dea, M. R. & Liddelow, S. A. Epigenetic memory astrocytes are likely an artifact of immune cell contamination. bioRxiv, 2025.2004.2004.647246 (2025). 10.1101/2025.04.04.647246

30 Mayo, L. et al. Regulation of astrocyte activation by glycolipids drives chronic CNS inflammation. Nat Med 20, 1147–1156 (2014). 10.1038/nm.3681

31 O’Shea, T. M. et al. Derivation and transcriptional reprogramming of border-forming wound repair astrocytes after spinal cord injury or stroke in mice. Nat Neurosci 27, 1505–1521 (2024). 10.1038/s41593-024-01684-6

32 Prakash, P. et al. Proteomic profiling of interferon-responsive reactive astrocytes in rodent and human. Glia 72, 625–642 (2024). 10.1002/glia.24494

33 Hasel, P., Rose, I. V. L., Sadick, J. S., Kim, R. D. & Liddelow, S. A. Neuroinflammatory astrocyte subtypes in the mouse brain. Nat Neurosci 24, 1475–1487 (2021). 10.1038/s41593-021-00905-6

34 Zamanian, J. L. et al. Genomic analysis of reactive astrogliosis. J Neurosci 32, 6391–6410 (2012). 10.1523/JNEUROSCI.6221-11.2012

35 Denstaedt, S. J. et al. S100A8/A9 Drives Neuroinflammatory Priming and Protects against Anxiety-like Behavior after Sepsis. J Immunol 200, 3188–3200 (2018). 10.4049/jimmunol.1700834

36 Burda, J. E. et al. Divergent transcriptional regulation of astrocyte reactivity across disorders. Nature 606, 557–564 (2022). 10.1038/s41586-022-04739-5

37 Fisher, T. M. & Liddelow, S. A. Emerging roles of astrocytes as immune effectors in the central nervous system. Trends Immunol 45, 824–836 (2024). 10.1016/j.it.2024.08.008

38 Guttenplan, K. A. et al. Knockout of reactive astrocyte activating factors slows disease progression in an ALS mouse model. Nat Commun 11, 3753 (2020). 10.1038/s41467-020-17514-9

39 Itoh, N. et al. Cell-specific and region-specific transcriptomics in the multiple sclerosis model: Focus on astrocytes. Proc Natl Acad Sci U S A 115, E302–E309 (2018). 10.1073/pnas.1716032115

40 Sardar, D. et al. Induction of astrocytic Slc22a3 regulates sensory processing through histone serotonylation. Science 380, eade0027 (2023). 10.1126/science.ade0027

41 Huang-Hobbs, E. et al. Remote neuronal activity drives glioma progression through SEMA4F. Nature 619, 844–850 (2023). 10.1038/s41586-023-06267-2

42 Yao, Z. et al. A high-resolution transcriptomic and spatial atlas of cell types in the whole mouse brain. Nature 624, 317–332 (2023). 10.1038/s41586-023-06812-z

43 Handsaker, R. E. et al. Long somatic DNA-repeat expansion drives neurodegeneration in Huntington’s disease. Cell 188, 623-639.e619 (2025). 10.1016/j.cell.2024.11.038

44 Falcão, A. M. et al. Disease-specific oligodendrocyte lineage cells arise in multiple sclerosis. Nat Med 24, 1837–1844 (2018). 10.1038/s41591-018-0236-y

45 Wohlfahrt, T. et al. PU.1 controls fibroblast polarization and tissue fibrosis. Nature 566, 344–349 (2019). 10.1038/s41586-019-0896-x

46 Kukanja, P. et al. Cellular architecture of evolving neuroinflammatory lesions and multiple sclerosis pathology. Cell 187, 1990–2009 e1919 (2024). 10.1016/j.cell.2024.02.030

47 Groh, A. M. R. et al. Ependymal cells undergo astrocyte-like reactivity in response to neuroinflammation. J Neurochem 168, 3449–3466 (2024). 10.1111/jnc.16120

48 Gruber, R. C. et al. BTK regulates microglial function and neuroinflammation in human stem cell models and mouse models of multiple sclerosis. Nat Commun 15, 10116 (2024). 10.1038/s41467-024-54430-8

49 Cameron, E. G. et al. A molecular switch for neuroprotective astrocyte reactivity. Nature 626, 574–582 (2024). 10.1038/s41586-023-06935-3

50 Kelley, K. W. et al. Kir4.1-Dependent Astrocyte-Fast Motor Neuron Interactions Are Required for Peak Strength. Neuron 98, 306–319 e307 (2018). 10.1016/j.neuron.2018.03.010

51 Irala, D. et al. Astrocyte-secreted neurocan controls inhibitory synapse formation and function. Neuron 112, 1657–1675 e1610 (2024). 10.1016/j.neuron.2024.03.007

52 Zhang, L. et al. Modulating mTOR-dependent astrocyte substate transitions to alleviate neurodegeneration. Nat Aging (2025). 10.1038/s43587-024-00792-z

53 Tao, X. et al. Astrocyte-conditional knockout of MOB2 inhibits the phenotypic conversion of reactive astrocytes from A1 to A2 following spinal cord injury in mice. Int J Biol Macromol 300, 140289 (2025). 10.1016/j.ijbiomac.2025.140289

54 Xu, J., Yan, Z., Bang, S., Velmeshev, D. & Ji, R. R. GPR37L1 identifies spinal cord astrocytes and protects neuropathic pain after nerve injury. Neuron (2025). 10.1016/j.neuron.2025.01.012

55 Tinkey, R. A., Smith, B. C., Habean, M. L. & Williams, J. L. BATF2 is a regulator of interferon-gamma signaling in astrocytes during neuroinflammation. Cell Rep 44, 115393 (2025). 10.1016/j.celrep.2025.115393

56 Wei, K. C., Lin, J. T. & Lin, C. H. Celecoxib paradoxically induces COX-2 expression and astrocyte activation through the ERK/JNK/AP-1 signaling pathway in the cerebral cortex of rats. Neurochem Int 183, 105926 (2025). 10.1016/j.neuint.2024.105926

57 Lin, C. H., Chen, H. Y. & Wei, K. C. Role of HMGB1/TLR4 Axis in Ischemia/Reperfusion-Impaired Extracellular Glutamate Clearance in Primary Astrocytes. Cells 9 (2020). 10.3390/cells9122585

58 Zarei-Kheirabadi, M., Mirsadeghi, S., Vaccaro, A. R., Rahimi-Movaghar, V. & Kiani, S. Protocol for purification and culture of astrocytes: useful not only in 2 days postnatal but also in adult rat brain. Mol Biol Rep 47, 1783–1794 (2020). 10.1007/s11033-020-05272-2

59 Chang, J. et al. Transplantation of A2 type astrocytes promotes neural repair and remyelination after spinal cord injury. Cell Commun Signal 21, 37 (2023). 10.1186/s12964-022-01036-6

60 Stogsdill, J. A. et al. Astrocytic neuroligins control astrocyte morphogenesis and synaptogenesis. Nature 551, 192–197 (2017). 10.1038/nature24638

61 Qin, L. et al. Astrocytic Neuroligin-3 influences gene expression and social behavior, but is dispensable for synapse number. Mol Psychiatry 30, 84–96 (2025). 10.1038/s41380-024-02659-6

62 Sun, M. et al. Leptin reduces LPS-induced A1 reactive astrocyte activation and inflammation via inhibiting p38-MAPK signaling pathway. Glia 73, 25–37 (2025). 10.1002/glia.24611

63 Meadows, S. M. et al. Hippocampal astrocytes induce sex-dimorphic effects on memory. Cell Rep 43, 114278 (2024). 10.1016/j.celrep.2024.114278

64 Orr, A. G. et al. Astrocytic adenosine receptor A2A and Gs-coupled signaling regulate memory. Nat Neurosci 18, 423–434 (2015). 10.1038/nn.3930

65 Baldwin, K. T. et al. HepaCAM controls astrocyte self-organization and coupling. Neuron 109, 2427–2442 e2410 (2021). 10.1016/j.neuron.2021.05.025

66 Jin, S. et al. Astroglial exosome HepaCAM signaling and ApoE antagonization coordinates early postnatal cortical pyramidal neuronal axon growth and dendritic spine formation. Nat Commun 14, 5150 (2023). 10.1038/s41467-023-40926-2

67 Takano, T. et al. Chemico-genetic discovery of astrocytic control of inhibition in vivo. Nature 588, 296–302 (2020). 10.1038/s41586-020-2926-0

68 Tang, W. et al. Calycosin regulates astrocyte reactivity and astrogliosis after spinal cord injury by targeting STAT3 phosphorylation. J Neuroimmunol 400, 578535 (2025). 10.1016/j.jneuroim.2025.578535

69 Molofsky, A. V. et al. Astrocyte-encoded positional cues maintain sensorimotor circuit integrity. Nature 509, 189–194 (2014). 10.1038/nature13161

70 Lopez-Hernandez, T. et al. Mutant GlialCAM causes megalencephalic leukoencephalopathy with subcortical cysts, benign familial macrocephaly, and macrocephaly with retardation and autism. Am J Hum Genet 88, 422–432 (2011). 10.1016/j.ajhg.2011.02.009

71 Taylor, X. et al. A1 reactive astrocytes and a loss of TREM2 are associated with an early stage of pathology in a mouse model of cerebral amyloid angiopathy. J Neuroinflammation 17, 223 (2020). 10.1186/s12974-020-01900-7

72 Windham, I. A. et al. APOE traffics to astrocyte lipid droplets and modulates triglyceride saturation and droplet size. J Cell Biol 223 (2024). 10.1083/jcb.202305003

73 Linnerbauer, M. et al. The astrocyte-produced growth factor HB-EGF limits autoimmune CNS pathology. Nat Immunol 25, 432–447 (2024). 10.1038/s41590-024-01756-6

74 Theparambil, S. M. et al. Adenosine signalling to astrocytes coordinates brain metabolism and function. Nature 632, 139–146 (2024). 10.1038/s41586-024-07611-w

75 Theparambil, S. M. et al. Astrocytes regulate brain extracellular pH via a neuronal activity-dependent bicarbonate shuttle. Nat Commun 11, 5073 (2020). 10.1038/s41467-020-18756-3

76 Turovsky, E. et al. Mechanisms of CO2/H+ Sensitivity of Astrocytes. J Neurosci 36, 10750–10758 (2016). 10.1523/JNEUROSCI.1281-16.2016

77 Wang, L. et al. Primary cilia signaling in astrocytes mediates development and regional-specific functional specification. Nat Neurosci 27, 1708–1720 (2024). 10.1038/s41593-024-01726-z

78 Yang, F. et al. Reactive astrocytes secrete the chaperone HSPB1 to mediate neuroprotection. Sci Adv 10, eadk9884 (2024). 10.1126/sciadv.adk9884

79 Kim, J. H. et al. Lipocalin-2 Is a Key Regulator of Neuroinflammation in Secondary Traumatic and Ischemic Brain Injury. Neurotherapeutics 20, 803–821 (2023). 10.1007/s13311-022-01333-5

80 Kim, J. H. et al. Aberrant activation of hippocampal astrocytes causes neuroinflammation and cognitive decline in mice. PLoS Biol 22, e3002687 (2024). 10.1371/journal.pbio.3002687

81 Liu, D. et al. Regulation of blood-brain barrier integrity by Dmp1-expressing astrocytes through mitochondrial transfer. Sci Adv 10, eadk2913 (2024). 10.1126/sciadv.adk2913

82 Pereira, A. et al. Direct neuronal reprogramming of mouse astrocytes is associated with multiscale epigenome remodeling and requires Yy1. Nat Neurosci 27, 1260–1273 (2024). 10.1038/s41593-024-01677-5

83 Russo, G. L. et al. CRISPR-Mediated Induction of Neuron-Enriched Mitochondrial Proteins Boosts Direct Glia-to-Neuron Conversion. Cell Stem Cell 28, 584 (2021). 10.1016/j.stem.2020.11.017

84 Zhou, X. et al. Pharmacological inhibition of Kir4.1 evokes rapid-onset antidepressant responses. Nat Chem Biol 20, 857–866 (2024). 10.1038/s41589-024-01555-y

85 Park, J. W. et al. Hypothalamic astrocyte NAD(+) salvage pathway mediates the coupling of dietary fat overconsumption in a mouse model of obesity. Nat Commun 15, 2102 (2024). 10.1038/s41467-024-46009-0

86 Ma, W. et al. Type I interferon response in astrocytes promotes brain metastasis by enhancing monocytic myeloid cell recruitment. Nat Commun 14, 2632 (2023). 10.1038/s41467-023-38252-8

87 Delgado, R. N. et al. Individual human cortical progenitors can produce excitatory and inhibitory neurons. Nature 601, 397–403 (2022). 10.1038/s41586-021-04230-7

88 Zhou, T. et al. Microglial debris is cleared by astrocytes via C4b-facilitated phagocytosis and degraded via RUBICON-dependent noncanonical autophagy in mice. Nat Commun 13, 6233 (2022). 10.1038/s41467-022-33932-3

89. Wu, Z. et al. A sensitive GRAB sensor for detecting extracellular ATP in vitro and in vivo. Neuron 110, 770-782 e775 (2022). 10.1016/j.neuron.2021.11.027

90 Monteiro, C. et al. Stratification of radiosensitive brain metastases based on an actionable S100A9/RAGE resistance mechanism. Nat Med 28, 752–765 (2022). 10.1038/s41591-022-01749-8

91 Smits, A. H. et al. Biological plasticity rescues target activity in CRISPR knock outs. Nat Methods 16, 1087–1093 (2019). 10.1038/s41592-019-0614-5

92 Squair, J. W. et al. Confronting false discoveries in single-cell differential expression. Nat Commun 12, 5692 (2021). 10.1038/s41467-021-25960-2

93 Rapaport, F. et al. Comprehensive evaluation of differential gene expression analysis methods for RNA-seq data. Genome Biol 14, R95 (2013). 10.1186/gb-2013-14-9-r95

94 Tarazona, S., Garcia-Alcalde, F., Dopazo, J., Ferrer, A. & Conesa, A. Differential expression in RNA-seq: a matter of depth. Genome Res 21, 2213–2223 (2011). 10.1101/gr.124321.111

95 Zhong, J. et al. Distinct roles of TREM2 in central nervous system cancers and peripheral cancers. Cancer Cell 42, 968–984 e969 (2024). 10.1016/j.ccell.2024.05.001

96 He, L. et al. CRISPR/Cas9/AAV9-mediated in vivo editing identifies MYC regulation of 3D genome in skeletal muscle stem cell. Stem Cell Reports 16, 2442–2458 (2021). 10.1016/j.stemcr.2021.08.011

97 Mountoufaris, G. et al. A line attractor encoding a persistent internal state requires neuropeptide signaling. Cell 187, 5998–6015 e5918 (2024). 10.1016/j.cell.2024.08.015

98 Mandl, M. et al. CRISPR/Cas9-mediated gene knockout in human adipose stem/progenitor cells. Adipocyte 9, 626–635 (2020). 10.1080/21623945.2020.1834230

99 Tuladhar, R. et al. CRISPR-Cas9-based mutagenesis frequently provokes on-target mRNA misregulation. Nat Commun 10, 4056 (2019). 10.1038/s41467-019-12028-5

100 Subramanian, A. et al. Gene set enrichment analysis: a knowledge-based approach for interpreting genome-wide expression profiles. Proc Natl Acad Sci U S A 102, 15545–15550 (2005). 10.1073/pnas.0506580102

101 Mews, P. et al. Alcohol metabolism contributes to brain histone acetylation. Nature 574, 717–721 (2019). 10.1038/s41586-019-1700-7

102 Schiroli, G. et al. Cell of origin epigenetic priming determines susceptibility to Tet2 mutation. Nat Commun 15, 4325 (2024). 10.1038/s41467-024-48508-6

103 Hochrein, S. M. et al. The glucose transporter GLUT3 controls T helper 17 cell responses through glycolytic-epigenetic reprogramming. Cell Metab 34, 516–532 e511 (2022). 10.1016/j.cmet.2022.02.015

104 Kang, T. G. et al. Epigenetic regulators of clonal hematopoiesis control CD8 T cell stemness during immunotherapy. Science 386, eadl4492 (2024). 10.1126/science.adl4492

105 Wheeler, M. A. et al. MAFG-driven astrocytes promote CNS inflammation. Nature 578, 593–599 (2020). 10.1038/s41586-020-1999-0

106 Sanmarco, L. M., Polonio, C. M., Wheeler, M. A. & Quintana, F. J. Functional immune cell-astrocyte interactions. J Exp Med 218 (2021). 10.1084/jem.20202715

107 Hasel, P. et al. Defining the molecular identity and morphology of glia limitans superficialis astrocytes in vertebrates. Cell Rep 44, 115344 (2025). 10.1016/j.celrep.2025.115344

108 de Ceglia, R. et al. Specialized astrocytes mediate glutamatergic gliotransmission in the CNS. Nature 622, 120–129 (2023). 10.1038/s41586-023-06502-w

109 Schirmer, L. et al. Neuronal vulnerability and multilineage diversity in multiple sclerosis. Nature 573, 75–82 (2019). 10.1038/s41586-019-1404-z

110 Absinta, M. et al. A lymphocyte-microglia-astrocyte axis in chronic active multiple sclerosis. Nature 597, 709–714 (2021). 10.1038/s41586-021-03892-7

111 Macnair, W. et al. snRNA-seq stratifies multiple sclerosis patients into distinct white matter glial responses. Neuron 113, 396–410 e399 (2025). 10.1016/j.neuron.2024.11.016

112 Gao, F. et al. Comparative analysis of three purification protocols for retinal ganglion cells from rat. Mol Vis 22, 387–400 (2016).

113 Zhang, P. W. et al. Generation of GFAP::GFP astrocyte reporter lines from human adult fibroblast-derived iPS cells using zinc-finger nuclease technology. Glia 64, 63–75 (2016). 10.1002/glia.22903

114 Krencik, R. & Zhang, S. C. Directed differentiation of functional astroglial subtypes from human pluripotent stem cells. Nat Protoc 6, 1710–1717 (2011). 10.1038/nprot.2011.405

115 Shaltouki, A., Peng, J., Liu, Q., Rao, M. S. & Zeng, X. Efficient generation of astrocytes from human pluripotent stem cells in defined conditions. Stem Cells 31, 941–952 (2013). 10.1002/stem.1334

116 Lafaille, F. G. et al. Impaired intrinsic immunity to HSV-1 in human iPSC-derived TLR3-deficient CNS cells. Nature 491, 769–773 (2012). 10.1038/nature11583

117 Serio, A. et al. Astrocyte pathology and the absence of non-cell autonomy in an induced pluripotent stem cell model of TDP-43 proteinopathy. Proc Natl Acad Sci U S A 110, 4697–4702 (2013). 10.1073/pnas.1300398110

118 Roybon, L. et al. Human stem cell-derived spinal cord astrocytes with defined mature or reactive phenotypes. Cell Rep 4, 1035–1048 (2013). 10.1016/j.celrep.2013.06.021

119 Chisholm, N. C. et al. Histone methylation patterns in astrocytes are influenced by age following ischemia. Epigenetics 10, 142–152 (2015). 10.1080/15592294.2014.1001219

120 Beurel, E. HDAC6 regulates LPS-tolerance in astrocytes. PLoS ONE 6, e25804 (2011). 10.1371/journal.pone.0025804

121 Pavlou, M. A. S. et al. Transcriptional and Chromatin Accessibility Profiling of Neural Stem Cells Differentiating into Astrocytes Reveal Dynamic Signatures Affected under Inflammatory Conditions. Cells 12 (2023). 10.3390/cells12060948

122 Wheeler, M. A. et al. Environmental Control of Astrocyte Pathogenic Activities in CNS Inflammation. Cell 176, 581–596 e518 (2019). 10.1016/j.cell.2018.12.012

123 Chao, C. C. et al. Metabolic Control of Astrocyte Pathogenic Activity via cPLA2-MAVS. Cell 179, 1483–1498 e1422 (2019). 10.1016/j.cell.2019.11.016

124 Clark, I. C. et al. Barcoded viral tracing of single-cell interactions in central nervous system inflammation. Science 372 (2021). 10.1126/science.abf1230

125 Rothhammer, V. et al. Microglial control of astrocytes in response to microbial metabolites. Nature 557, 724–728 (2018). 10.1038/s41586-018-0119-x

126 Sanmarco, L. M. et al. Gut-licensed IFNγ(+) NK cells drive LAMP1(+)TRAIL(+) anti-inflammatory astrocytes. Nature 590, 473–479 (2021). 10.1038/s41586-020-03116-4

127 Foo, L. C. Purification of rat and mouse astrocytes by immunopanning. Cold Spring Harb Protoc 2013, 421–432 (2013). 10.1101/pdb.prot074211

128 Gutierrez-Vazquez, C. & Quintana, F. J. Protocol for in vitro analysis of pro-inflammatory and metabolic functions of cultured primary murine astrocytes. STAR Protoc 3, 101033 (2022). 10.1016/j.xpro.2021.101033

129 Foo, L. C. et al. Development of a method for the purification and culture of rodent astrocytes. Neuron 71, 799–811 (2011). 10.1016/j.neuron.2011.07.022

130 Zhang, Y. et al. Purification and Characterization of Progenitor and Mature Human Astrocytes Reveals Transcriptional and Functional Differences with Mouse. Neuron 89, 37–53 (2016). 10.1016/j.neuron.2015.11.013

131 Tchieu, J. et al. NFIA is a gliogenic switch enabling rapid derivation of functional human astrocytes from pluripotent stem cells. Nature Biotechnology 37, 267–275 (2019). 10.1038/s41587-019-0035-0

132 Robinson, M. D., McCarthy, D. J. & Smyth, G. K. edgeR: a Bioconductor package for differential expression analysis of digital gene expression data. Bioinformatics 26, 139–140 (2010). 10.1093/bioinformatics/btp616

133 Gu, Z. Complex heatmap visualization. Imeta 1, e43 (2022). 10.1002/imt2.43

134 Love, M. I., Huber, W. & Anders, S. Moderated estimation of fold change and dispersion for RNA-seq data with DESeq2. Genome Biol 15, 550 (2014). 10.1186/s13059-014-0550-8

135 McGinnis, C. S., Murrow, L. M. & Gartner, Z. J. DoubletFinder: Doublet Detection in Single-Cell RNA Sequencing Data Using Artificial Nearest Neighbors. Cell Syst 8, 329–337 e324 (2019). 10.1016/j.cels.2019.03.003

136 Korsunsky, I. et al. Fast, sensitive and accurate integration of single-cell data with Harmony. Nat Methods 16, 1289–1296 (2019). 10.1038/s41592-019-0619-0

137 Satija, R., Farrell, J. A., Gennert, D., Schier, A. F. & Regev, A. Spatial reconstruction of single-cell gene expression data. Nat Biotechnol 33, 495–502 (2015). 10.1038/nbt.3192

138 Yang, Q. et al. PANoptosis, an indicator of COVID-19 severity and outcomes. Brief Bioinform 25 (2024). 10.1093/bib/bbae124

139 Liu, J. et al. Autophagy-prominent cell clusters among human lens epithelial cells: integrated single-cell RNA-sequencing analysis. BMC Ophthalmol 23, 168 (2023). 10.1186/s12886-023-02910-8

140 Dai, S. L., Pan, J. Q. & Su, Z. R. Multi-omics features of immunogenic cell death in gastric cancer identified by combining single-cell sequencing analysis and machine learning. Sci Rep 14, 21751 (2024). 10.1038/s41598-024-73071-x

141 Tirosh, I. et al. Dissecting the multicellular ecosystem of metastatic melanoma by single-cell RNA-seq. Science 352, 189–196 (2016). 10.1126/science.aad0501

